# High throughput screening of mesenchymal stromal cell morphological response to inflammatory signals for bioreactor-based manufacturing of extracellular vesicles that modulate microglia

**DOI:** 10.1101/2023.11.19.567730

**Authors:** Andrew M. Larey, Thomas M. Spoerer, Kanupriya R. Daga, Maria G. Morfin, Hannah M. Hynds, Jana Carpenter, Kelly M. Hines, Ross A. Marklein

## Abstract

Due to their immunomodulatory function, mesenchymal stromal cells (MSCs) are a promising therapeutic with the potential to treat neuroinflammation associated with neurodegenerative diseases. This function can be mediated by secreted extracellular vesicles (MSC-EVs). Despite established safety, MSC clinical translation has been unsuccessful due to inconsistent clinical outcomes resulting from functional heterogeneity. Current approaches to mitigate functional heterogeneity include ‘priming’ MSCs with inflammatory signals to enhance function. However, comprehensive evaluation of priming and its effects on MSC-EV function has not been performed. Clinical translation of MSC-EV therapies requires significant manufacturing scale-up, yet few studies have investigated the effects of priming in bioreactors. As MSC morphology has been shown to predict their immunomodulatory function, we screened MSC morphological response to an array of priming signals and evaluated MSC-EV identity and potency in response to priming in flasks and bioreactors. We identified unique priming conditions corresponding to distinct morphologies. These conditions demonstrated a range of MSC-EV preparation quality and lipidome, allowing us to discover a novel MSC-EV manufacturing condition, as well as gain insight into potential mechanisms of MSC-EV microglia modulation. Our novel screening approach and application of priming to MSC-EV bioreactor manufacturing informs refinement of larger-scale manufacturing and enhancement of MSC-EV function.

**Graphical Abstract:** 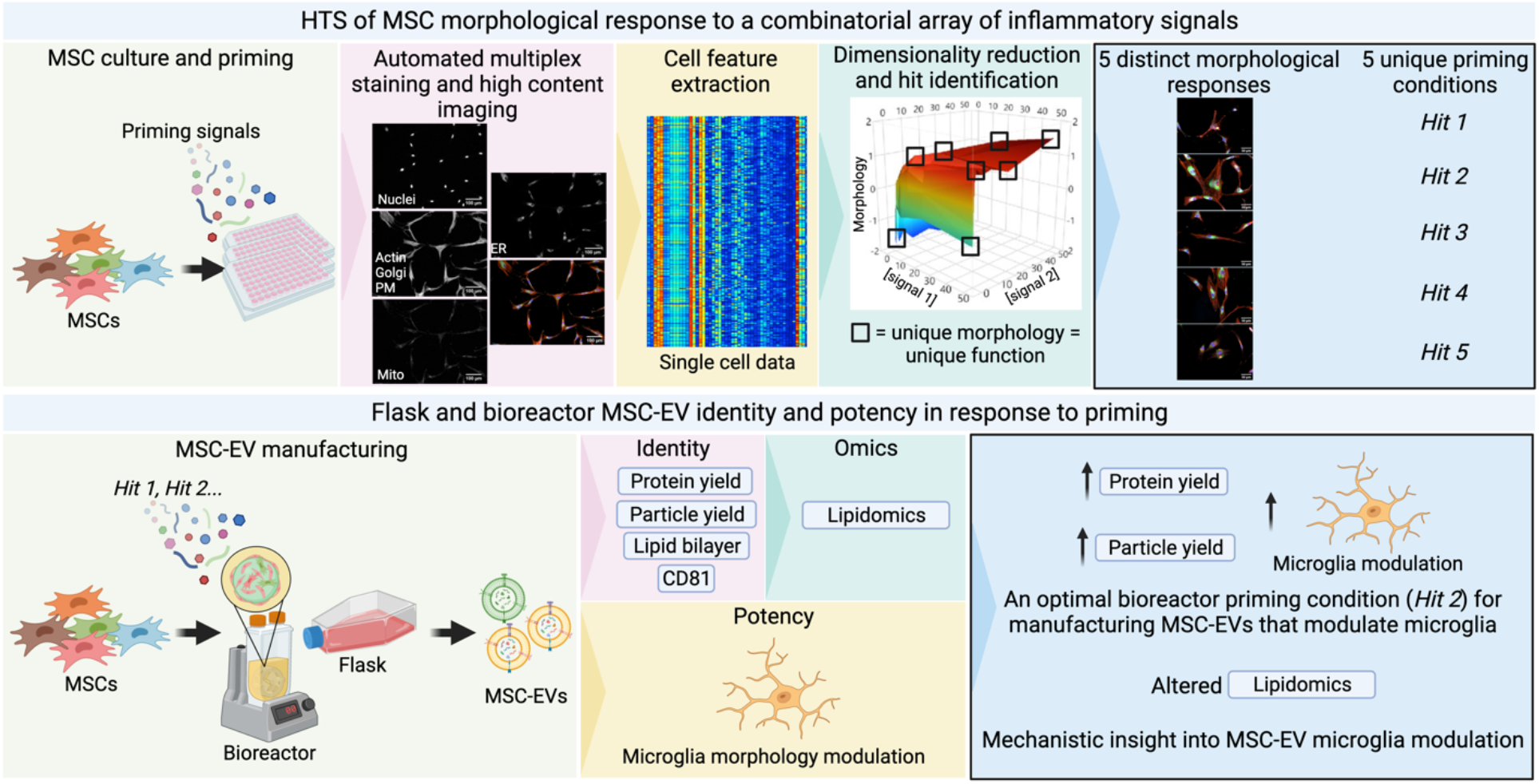

**Highlights:**

- MSCs morphologically respond to inflammatory priming conditions.
- Priming ‘hits’ identified from morphological screen increase MSC-EV production.
- Priming MSCs in bioreactors enhances MSC-EV modulation of microglia.
- Changes in MSC-EV production and potency reflected by lipid content.
- First demonstration of effects of priming on MSC production of EVs in a bioreactor.

## INTRODUCTION

Mesenchymal stromal cells (MSCs) are a promising cell therapy due to their established immunomodulatory and regenerative effects (1, 2). MSCs can be derived from various tissue sources including bone marrow, adipose tissue, and umbilical cord blood and are often characterized based on the International Society of Cell and Gene Therapy (ISCT) criteria of 1) plastic adherence, 2) surface marker phenotype, and 3) trilineage differentiation capacity (osteocytes, chondrocytes, and adipocytes) (3). However, it has become increasingly recognized that MSC therapeutically relevant functions are mediated through paracrine signaling (4). MSC immunomodulation has relevance for treating diseases with overactive immune components including Alzheimer’s Disease (5), multiple sclerosis (6), traumatic brain injury (7), and acute respiratory distress syndrome (8). MSC immunomodulatory function can be mediated by secreted extracellular vesicles (MSC-EVs), which are lipid-bilayer enclosed, highly regulated signaling vehicles containing a host of bioactive factors such as proteins, lipids, and RNAs (9, 10). Compared to the MSCs that produce them, MSC-EVs could be a more promising therapeutic as they have a lower risk of tumorigenicity (11), can readily cross the blood-brain barrier (12), and can potentially be stored more economically (i.e., at room temperature) (13).

Despite a well-established safety profile across approximately 1,100 clinical trials and a wealth of data showing *in vitro* and *in vivo* efficacy for numerous regenerative medicine applications (14, 15), MSC clinical translation has been generally unsuccessful (i.e., no FDA approved MSC-based therapies) due to inconsistent clinical outcomes resulting from functional heterogeneity (16). MSC functional heterogeneity is impacted by material (e.g., tissue sources, subpopulations, culture reagents) and process (e.g., platform, scale, automation) parameters (17). Numerous studies have demonstrated significant donor-dependent differences in MSC identity and potency (18–21). Current approaches to mitigating functional heterogeneity include stimulating MSCs with physiologically relevant inflammatory conditions (termed ‘priming’), which enhances immunomodulatory function (22). Common priming strategies involve stimulation with cytokines such as IFN-γ and TNF-α or hypoxia (0.5-5% [O2]) transiently in culture (23). Furthermore, changes in culture geometry (e.g., 2D flask culture to 3D hydrogel culture) show significant enhancement of regenerative and immunomodulatory functions (24, 25). Primed MSCs demonstrate enhanced immunomodulatory function compared to unprimed MSCs targeting T cells (reduced CD4+ and CD8+ T cell proliferation (26–33); T_reg_ cell induction (34–36)), NK cells (reduced proliferation (37)), B cells (reduced proliferation (37)), neutrophils (activation and recruitment (38)), and macrophages (reduced cytokine secretion (39)). Similar to the producing MSCs, MSC-EV immunomodulatory function has been shown to improve in response to priming with target cells of macrophages (M1 to M2 polarization (40–48)), T cells (reduced proliferation, reduced cytokine secretion (29, 49, 50); T_reg_ cell induction (51–53)), NK cells (reduced proliferation (37)), and B cells (reduced proliferation (37))(54). However, a systematic exploration of combinatorial priming cues and the effects on MSC-EV immunomodulatory function has not been conducted.

Clinical translation (and eventual commercialization) require significant scale-up of MSC manufacturing to achieve EV production in accordance with dose, number of patients, and number of administrations per year for a given disease application (55, 56). Optimal priming conditions for a given application must be identified prior to clinical scale manufacturing where screening of many conditions is prohibitively expensive. Moreover, necessary changes in manufacturing format (e.g., flasks to stirred tank, wave bed, or hollow fiber bioreactors (57)) and scale (e.g., from <1 L to 1E1–1E3 L) in the progression of therapies from preclinical to early (Phase I) and late stage (Phase 3) clinical trials can have an impact on product quality (i.e., introduce heterogeneity) and necessitate comparability studies (58–60). However, very few studies have explored the effects of priming in larger-scale bioreactors. Furthermore, evaluating the effects of priming in bioreactors entails understanding of the fundamental regulation of the MSC and EV functional response to priming in larger-scale formats and effective prediction of this response.

We have established MSC morphology as a putative CQA (critical quality attribute, predictor of function) for immunomodulation (26, 61, 62). Cell morphology can be measured in a low-cost, high-throughput manner and provides a visual high-dimensional readout (‘morphological profile’) that reflects complex intracellular state (63, 64). Therefore, the objective of this study was to perform a high throughput screen of MSC morphological response to a combinatorial array of priming signals and to evaluate MSC-EV identity and immunomodulatory potency in response to select priming conditions in flasks and larger-scale bioreactor manufacturing formats. We hypothesized that our morphological profiling approach would identify unique priming conditions corresponding to unique MSC morphologies and MSC-EV function. Our focused screen of inflammatory-relevant signals identified unique priming condition ‘hits’ corresponding to unique morphologies. Select priming hits demonstrated a range of MSC-EV preparation quality, allowing us to discover novel, improved MSC-EV manufacturing conditions, as well as provide insight into potential metabolic regulators of MSC-EV functional response to priming.

## MATERIALS AND METHODS

### MSC Morphology and Cell Painting

Human adipose derived MSC line RB62 (ADMSC RB62) (RoosterBio) was expanded according to recommended RoosterBio protocol described previously (26) and then cryopreserved as passage 2 (P2). Cell line information for all MSC cell lines used in this study is in **Table S1**. At the time of the assays, ADMSC RB62 P2 was thawed into T175 flasks at a seeding density of 5,714 cells/cm^2^ and recovered in MSC growth media (MSC-GM) consisting of 10% FBS (Neuromics), 1% L-glutamine (Gibco), and 1% penicillin/streptomycin (Gibco) in alpha-MEM (Gibco). MSCs were cultured to 80% confluence following standard protocols (65). Viability after recovery and expansion was verified >90% for all assays by AO/PI staining (Nexcelom) and automated cell counting (Nexcelom Cellometer K2). MSCs were then passaged into 96-well plates at a seeding density of 1,562 cells/cm^2^ (n=8 wells in 4 replicate plates) and cultured for 24 hours. MSCs were treated with either MSC-GM or MSC-GM + 50 ng/mL IFN-γ/TNF-α (Gibco, Sino Biological, respectively) for 24 hours. Morphology fixing and staining was performed by treating MSCs with 4% paraformaldehyde (Electron Microscopy Sciences) for 15 minutes, washing 1X with PBS −/− (Corning), treating with 10 ug/mL Hoechst nuclear stain (Invitrogen) + 20 uM fluorescein-5-maleimide cytoplasm stain (ThermoFisher) + 0.1% (v/v) Tween-20 (Sigma-Aldrich) for 1 hour, and washing 3X with PBS −/−. Modified Cell Painting (66) fixing and staining was performed by treating MSCs with 10 ng/mL MitoTracker Deep Red mitochondria stain (Invitrogen) for 30 minutes at 37° C, fixing with 4% paraformaldehyde for 30 minutes, washing 2X with HBSS (Gibco), then staining with 1% (w/v) bovine serum albumin (Sigma-Aldrich) HBSS + 0.1% TritonX-100 (Sigma-Aldrich) + 1.5 μg/mL Wheat Germ Agglutinin Alexa Fluor 555 Conjugate plasma membrane stain (Invitrogen) + 8.25 nM Phalloidin/AlexaFluor 568 conjugate F actin cytoskeleton stain (Invitrogen) + 10 µg/mL Hoechst nuclei stain (Invitrogen) + 5 µg/mL concanavalin A/Alexa Fluor 488 conjugate endoplasmic reticulum stain (Invitrogen) for 30 minutes. Next, MSCs were washed 3X with HBSS. All morphology and Cell Painting liquid handling (fixing, staining, washing) was performed using a BioTek MultiFlo FX automated liquid handler (Agilent).

Each morphology well was imaged at 10X magnification in GFP and DAPI channels using a 6×6 non-overlapping montage (total area imaged/well = 0.173 cm^2^) and each Cell Painting well was imaged following the same specifications in GFP, DAPI, TR, and CY5 channels. All high content imaging (HCI) was performed using a BioTek Cytation5 automated microscope (Agilent). Morphological feature quantification was performed using custom morphology (**S1**) and modified Cell Painting (**S2**) CellProfiler (67) pipelines to generate high dimensional single cell data. Well-medians of every feature were calculated using a custom Python script (**S3**) and the mean of these well medians plotted for analysis of individual features. For composite morphological features, principal component analysis (PCA) was conducted on correlations in JMP Pro 16 using 21 morphological features based on previous work (62). The CTL-primed differential was calculated for every feature as

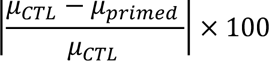

### Exploratory HTS

ADMSC RB62 P2 was thawed and recovered as described above using xeno-free MSC expansion media (RoosterNourish-MSC-XF, RoosterBio). MSCs were passaged into 96-well plates at a seeding density of 1,562 cells/cm^2^ and allowed to adhere overnight. MSCs were then treated with IFN-γ, TNF-α, and IL1-β at 0, 2, 10, and 50 ng/mL; ph 7.4, 6.8, and 6.2; and 20% (normoxia) and 2% (hypoxia) [O2] in a full factorial experimental design (384 total priming conditions, n=4 wells/priming condition) along with −CTL conditions with no cytokines, 7.4 pH, and 20% [O2] (n=12 wells/plate) and +CTL conditions with 50 ng/mL IFN-γ/TNF-α, 7.4 pH, and 20% [O2] (n=4 wells/plate) for 24 hours. All treatments were performed in an EV collection media (RoosterCollect-EV, RoosterBio). Design of experiments (DOE) in JMP Pro 16 was used to create semi-random plate layouts consisting of randomized quadrants of unique conditions repeated in a plate for 4 replicate wells of each condition. Modified Cell Painting and HCI was performed as described above with randomization of plate processing order to control for potential batch effects. Morphological feature quantification was performed using modified open source Cell Painting CellProfiler pipelines and Python scripts (Cell Painting GitHub, PyCytominer GitHub) (68, 69) on high performance computing (HPC) (Georgia Advanced Computing Resource Center, University of Georgia). First, illumination correction was performed for every image (S4). Next, morphological feature quantification was performed (**S2**) to generate high-dimensional single cell data. Modified Cell Painting Python scripts (69, 70) were then used to 1) aggregate all features to well medians (**S5**), 2) normalize all features to on-plate −CTLs using the formula (**S5**)

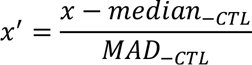

where MAD is median absolute deviation (**S5**), and 3) feature select using Pearson’s correlation coefficient to remove features with a correlation coefficient >0.9 (**S5**). Then, PCA was performed as described above.

For hit identification, the number of principal components (PCs) required to summarize 90% of the variance in the data were used in a Python script (Cytominer-eval GitHub) (70) (**S6**) to perform multidimensional perturbation value testing (mp-value testing) (71) to identify wells significantly different from the −CTL. PCA of significant mp-value wells was performed and the number of principal components required to summarize 90% of the variance in the data was used in Ward’s hierarchical clustering of significant mp-value wells. Priming conditions with all 4 replicate wells in the same cluster (an indication of low variability conditions) were considered to identify the most representative priming condition in a cluster, resulting in 10 hit priming conditions from the exploratory high throughput screen (HTS).

### Validation HTS

ADMSC RB62, bone marrow derived MSC line RB71 (BMMSC RB71) (RoosterBio), and adipose derived MSC line (ADMSC RB98) (RoosterBio) (expanded and cryopreserved at P2 as described above) were thawed and recovered as described for the exploratory HTS. The validation HTS was performed identically to the exploratory HTS through PCA and prior to hit identification using these 3 MSC lines and the 10 hits identified in the exploratory HTS.

### Exploratory MSC-EV manufacturing

Flask and bioreactor MSC-EV manufacturing: BMMSC RB71 P2 was thawed into T225 flasks at a seeding density of 4,444 cells/cm^2^ and recovered in RoosterNourish-MSC-XF through culture to 80% confluence following standard protocols. MSCs were then passaged into T175 flasks at a seeding density of 714 cell/cm^2^ or 0.5 L spinning wheel bioreactors (PBS Biotech) with 0.4 g Corning Synthemax II microcarriers for parallel flask and bioreactor culture. MSCs were expanded in flasks and bioreactors according to the standard protocols from PBS Biotech and RoosterBio. After expansion, MSCs in flasks and bioreactors were primed in RoosterCollect-EV with control and priming hits identified from the morphological screen.

MSC-EV preparation: MSC-EVs were prepared according to a modified 2-step ultracentrifugation (UC) protocol (72). Whole conditioned media from flasks and bioreactors was 0.2 μm filtered prior to first round UC at an RCFmax of 133,900 x g (Sorvall WX ultraCentrifuge, ThermoFisher; Fiberlite F37L-8×100 Fixed-Angle Rotor, ThermoFisher, k factor=168; PC Bottle Assembly 70 mL, ThermoFisher) for 1 hour at 4° C. The pellet was resuspended in ice cold PBS +/+ (Corning) for second round UC and transferred into microultracentrifuge tubes (PC Thickwall 4 mL, ThermoFisher) for microultracentrifugation at an RCFmax of 140,000 x g (Sorvall MX 150+ Micro-Ultracentrifuge, ThermoFisher; S110-AT Fixed-Angle Rotor, ThermoFisher, k factor=15.4) for 1 hour at 4° C. Pellets were finally resuspended in ice cold PBS +/+ to achieve a 37.5X concentration of the 0.20 μm filtered conditioned media processed in the first round UC. All MSC-EV preparations were stored at −80° C for <1 week, thawed, vortexed for 5 seconds, and quality assessment performed.

### EV Preparation Quality Assessment

Protein yield: MSC-EV preparation total protein concentration was determined using the Micro BCA Protein Assay Kit (ThermoFisher) according to the manufacturer’s protocol. Briefly, 75 µL of MSC-EV preparation was combined with 75 µL of the reagent mixture in a 96-well plate, uniformly mixed, and incubated for 2 hours at 37° C. Absorbance was read at 562 nm using a SpectraMax iD5 Multi-Mode Microplate Reader (Molecular Devices) and total protein concentration of the MSC-EV preparations was determined from the standard curve.

Particle yield: MSC-EV preparation particle concentration was determined using microfluidic tunable resistive pulse sensing on a Spectradyne nCS1 (Spectradyne). First, 200 nm polystyrene bead standards at a concentration of 1E9 beads/mL were counted to rescale diameter and concentration during data analysis. MSC-EV preparations were diluted 2X in dilution buffer for counting. Bead standards and all MSC-EV preparations were counted for 3 technical replicates using 3 C-400 cartridges and recording >1000 events with >75 nm diameter. Default cartridge peak filters of signal/noise and transit time as well as custom filters of diameter (75-400 nm) and symmetry (0.2-100,000) were applied to all replicates in Viewer software (Spectradyne) and resultant particle concentrations and median diameters exported.

Phenotype: MSC-EV preparation vesicular structure and canonical surface marker expression were determined using EV bead-based flow cytometry with human CD81 detection beads (Invitrogen) according to the manufacturer’s protocol. Briefly, beads were washed with buffer using a magnetic separator prior to addition of MSC-EV preparations and incubation at 4° C overnight on an orbital shaker. Next, beads+MSC-EV preparation were washed and stained with 40 µM CFSE (BioLegend) for 1 hour at room temperature. Beads+MSC-EV preparation were then washed 2X, resuspended, and transferred to a round bottom 96 well plate for flow cytometry. Flow cytometry of bead bound EVs was performed using a Quanteon (Agilent) with >10,000 events collected per sample. Flow cytometry data was analyzed using FlowJo (Tree Star). Briefly, single beads were gated using scatter principles. Then, unstained controls were used to determine CFSE+ populations. The gating strategy is shown in **Fig. S1**.

Morphology-based microglia modulation assay: C20s, an immortalized human cell line, were generously provided by Drs. Min-Ho Kim and Eric Dyne (Kent State University). The growth media consisted of DMEM-F12 1:1 (Gibco), 10% FBS (Neuromics), 1% penicillin/streptomycin (Gibco), and 0.2% normocin (InvivoGen). C20s were thawed in growth media containing 1% N2 supplement (Gibco) for efficient recovery and 1 µM dexamethasone (Millipore Sigma). The cells were recovered post thaw for 48 hours, reaching 90% confluence for harvesting and seeding for experiments. For morphology assays, C20s were seeded in a Poly-D lysine hydrobromide (PDL) (Sigma-Aldrich) coated 96-well plate (Corning; Greiner Bio-One) at a seeding density of 1,500 cells/cm^2^ and allowed to adhere overnight. Then, cells were stimulated using a half media change with a final cytokine cocktail of 5 ng/mL IFN-ү/TNF-α (Gibco, Sino Biological, respectively). MSC-EV treatment groups received an additional 10 uL of MSC-EV preparation for the 1X dilution. All non-EV treatment controls were treated with 10 uL PBS +/+ only. After 24 hours, cells were fixed with 4% paraformaldehyde (Electron Microscopy Sciences) for 30 minutes, washed 2X with HBSS (Gibco), then stained with 1% (w/v) bovine serum albumin (Sigma-Aldrich) HBSS + 0.1% TritonX-100 (Sigma-Aldrich) + 1.5 μg/mL Wheat Germ Agglutinin Alexa Fluor 555 Conjugate plasma membrane stain (Invitrogen) + 8.25 nM Phalloidin/AlexaFluor 568 conjugate F actin cytoskeleton stain (Invitrogen) + 10 µg/mL Hoechst nuclei stain (Invitrogen) for 30 minutes. Next, C20s were washed 3X with HBSS. The 96-well plates were imaged using a 10X objective on the BioTek Cytation5 (Agilent) automated microscope. 36 fields of view were captured per well. Morphological feature quantification was performed using a custom CellProfiler pipeline to generate high dimensional single cell data. Well-medians of every feature were calculated using a custom Python script (**S3**) and normalized to on-plate unstimulated/stimulated microglia controls according to the formula

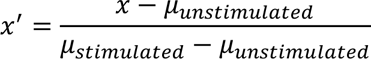

Composite microglia morphology PC1 score was calculated from PCA on correlations in JMP Pro 16 using 21 morphological features based on previous work (62). Batch-batch normalization was performed by subtracting the difference between −CTL1 and −CTL2 from −CTL2 within a scale (aligning −CTLs between batches) and subtracting the same difference from all batch 2 MSC-EV treatments.

MSC-EV preparation quality assessments are summarized in **Table S2**.

### MSC-EV Preparation Lipidomics

Lipid Extraction: Collected cells were washed with PBS and aliquoted into three-400 µL replicates before using a Bligh and Dyer (B&D) lipid extraction protocol (73). An additional 400 µL of the cell culture media was extracted directly for comparison. Washed cells received an additional 100 µL of HPLC grade H2O and were sonicated for 30 minutes at 4° C. We added 2 mL of a chilled solution of 1:2 CHCl3/MeOH to the sample and vortexed for 5 minutes. After, we added 0.5 mL of chilled CHCl3 and 0.5 mL of chilled H2O to facilitate phase separation. We then vortexed the samples for an additional 1 minute before centrifuging for 10 minutes at 3,500 rpm at 4° C. The lower organic layer of the biphasic solution was collected into clean glass tubes and dried under vacuum. Cell and culture media extracts were then reconstituted in 250 µL of 1:1 CHCl3/MeOH and stored at −-80° C.

Sample Preparation: Extracts were allowed to thaw to room temperature and were then prepared as 2X diluted (cell extracts) and 3-fold concentrated (culture media) samples in 95:5 ACN/H2O, 10 mM ammonium acetate (HILIC A) for LC-MS analysis. Cell extracts were also prepared as 1.5-2X dilutions in HILIC A for LC-MS/MS as well.

HILIC Chromatography: The lipid extracts were analyzed by hydrophilic interaction chromatography (HILIC) coupled to ion mobility-mass spectrometry. The HILIC separation was performed on a Waters Acquity FTN I-Class Plus ultraperformance liquid chromatography system using a Waters AQUITY UPLC BEH HILIC column (2.1 x 100 mm, 1.7 μm) maintained at 40° C. The mobile phases consisted of 95:5 ACN/H2O, 10 mM ammonium acetate (HILIC A, B) and 50:50 ACN/H2O, 10 mM ammonium acetate (HILIC B, A). A flow rate of 0.5 mL/min was used with the following gradient elution conditions: 0-0.5 min, 100% B; 0.5-5 min, ramp to 60% B; 5-5.5 min, 60% B; 5.5-6 min, ramp to 100% B; 6-7 min, 100% B. The autosampler chamber was maintained at 6° C. An injection volume of 10 μL was used.

Mass Spectrometry Analysis: Data were collected on a Waters SYNAPT XS traveling-wave ion mobility-mass spectrometer (TWIM-MS) in both positive and negative electrospray ionization mode with the following source conditions: capillary voltage, 2.0 kV (neg) and 3.0 kV (pos); sampling cone voltage, 40 V; source offset, 4 V; source temperature, 150⁰ C; desolvation temperature, 500⁰ C; desolvation gas flow rate, 1000 L/h; cone gas flow rate, 50 L/h. TWIM separations were performed in nitrogen with a gas flow of 90 mL/min, a wave velocity of 550 m/s, and a wave height of 40 V. Mass calibration was performed with sodium formate over the range of 50-1200 m/z. The time-of-flight mass analyzer was operated in V-mode (resolution mode) with a resolution of ∼30,000. Data were collected with a 1 s scan time over the range of 50-1200 m/z. Leucine enkephalin was used for continuous lock-mass correction during acquisition. For HILIC-IM-MS, MS/MS spectra were acquired using data-independent acquisition (MSe) with a ramped collision energy, 35 to 50 eV (pos) and 40 to 50 eV (neg), in the transfer region of the instrument. For HILIC-IM-MS/MS, precursors were selected in the quadrupole with an LM resolution of 12 and spectra were acquired with a ramped collision energy of 45 to 60 eV in the transfer region of the instrument.

### Statistical Analysis

All statistical tests were performed in GraphPad Prism 9 with the specific tests utilized for each experiment described in the figure legends.

## RESULTS

### Cell Painting enabled comprehensive, unbiased assessment of consistent MSC morphological response to cytokine priming

ADMSC RB62 was expanded and seeded into 96-well plates before treatment with MSC-GM or MSC-GM + 50 ng/mL IFN-γ/TNF-α. Morphological response to priming was quantified using a custom CellProfiler (67) pipeline to establish the morphology assay. MSC proliferation did not significantly change with priming such that cell count was consistent across wells and plates within an experiment, indicating consistent cell seeding (**Fig. 1Ai**). Three exemplary individual features encompassed overall changes in cell size, elongation, and complexity (cell area, cell aspect ratio, and cell form factor, respectively). Primed MSCs were larger (increased cell area), less elongated (decreased aspect ratio), and more round (higher form factor) compared to unprimed MSCs controls (**Fig. 1Aii-iv**). PCA of 21 features (selected based on previous work (62)) was used to understand changes in high dimensional morphological profile where the dominant source of variability (separated on PC1) was priming condition, rather than plate (**Fig. 1Av**). **Fig. 1B** illustrates representative cells stained with the morphological profiling approach previously used by our group (26, 61, 62) that only assesses overall cell and nuclear morphology. A modified Cell Painting (66) protocol was tested to improve discrimination of control and primed groups through 5-plex staining in 4 channels (Hoechst: nuclei; phalloidin/wheat germ agglutinin: actin fibers/Golgi/plasma membrane; MitoTracker: mitochondria; concanavalin: endoplasmic reticulum) (**Fig. 1D**) as opposed to 2-plex staining in 2 channels in the morphology assay (Hoechst: nuclei; fluorescein-5-maleimide: cytoplasm). Addition of intensity and texture features made the control-primed differential significantly higher in Cell Painting compared to morphology (**Fig. 1C**). Because this increased dynamic range (reflected by a greater number of features with high differential) could better enable hit identification, Cell Painting was used as an improved screening tool over the previous morphology assay.

**Figure 1.**
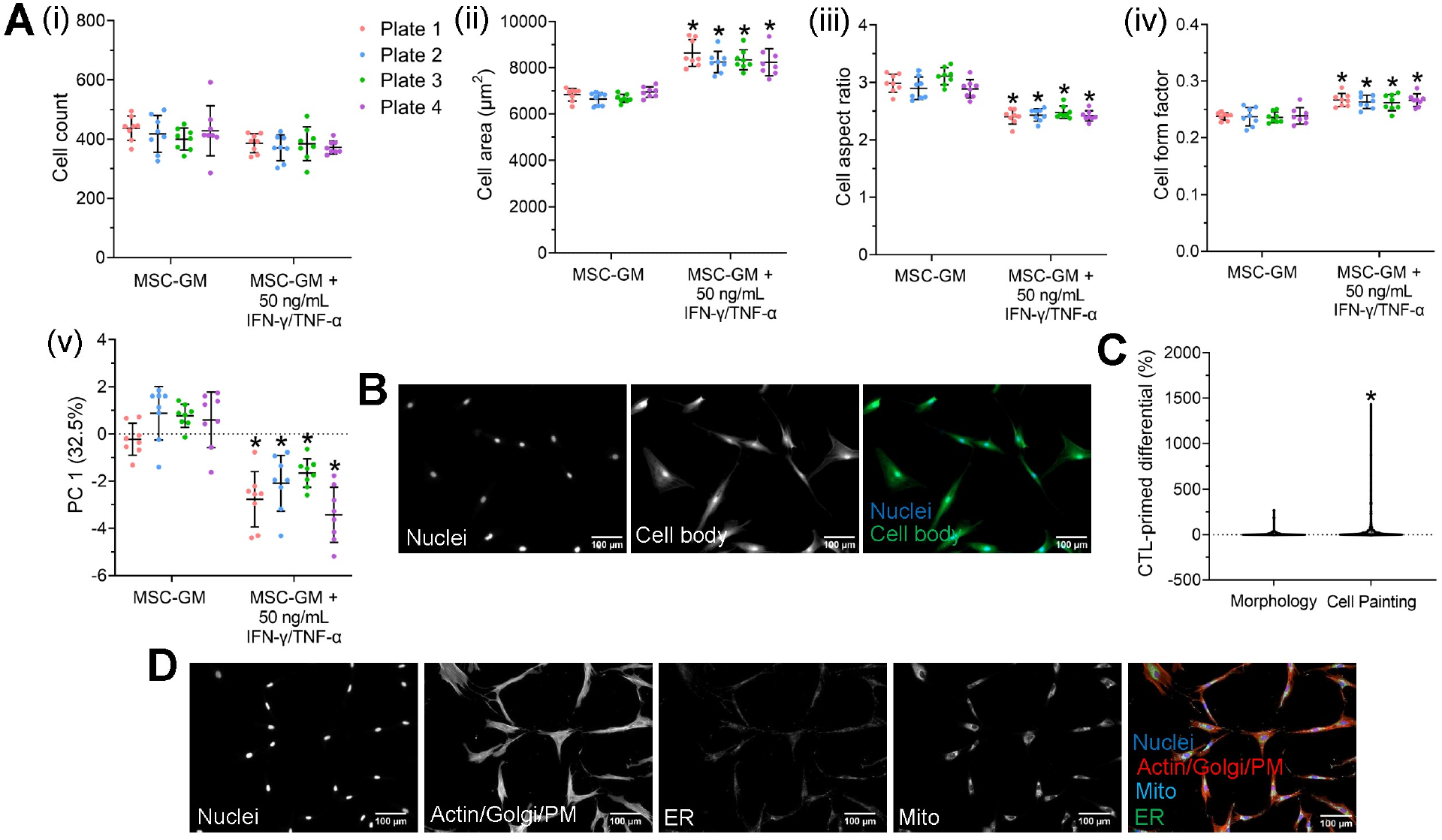
Cell Painting enabled comprehensive, unbiased assessment of consistent MSC morphological response to cytokine priming. **Ai.** Morphology assay cell count. **Aii-iv.** Morphology assay, three exemplary individual features. **Av.** Morphology assay composite morphological profile using PCA (input 21 features). **Ai-v.** Graphed as mean and standard deviation where each point represents 1 well (n = 8), *p<0.05 vs MSC-GM group within a plate, ordinary 2-way ANOVA with Šidak’s multiple comparisons test. **B.** Morphology representative cells, greyscale and color composite images. **C.** MSC-GM—MSC-GM + 50 ng/mL IFN-γ/TNF-α differential, Morphology (n = 113 features), Cell Painting (n = 566 features), *p<0.05 vs Morphology, Mann Whitney test. **D.** Cell Painting, representative cells greyscale and color composite images. Abbreviations: PM: plasma membrane; ER: endoplasmic reticulum; Mito: mitochondria.

### The exploratory HTS identified unique priming conditions corresponding to distinct morphological responses

A robust HTS and hit priming condition identification process was employed to discover unique priming conditions that we hypothesized correspond to unique MSC-EV function. **Fig. S2** outlines the experimental design, including priming conditions tested: IFN-γ, TNF-α, and IL-1β at 0, 2, 10, and 50 ng/mL; ph 7.4, 6.8, and 6.2; and 20% (normoxia) and 2% (hypoxia) [O2] in a full factorial experimental design (384 total priming conditions). These conditions and doses were selected because they are well-established inflammation relevant signals (39, 74–80). The +CTL (50 ng/mL IFN-γ + 50 ng/mL TNF-α) was selected based on past results of a synergistic priming effect contributing to a significant morphological response in multiple cell-lines (26). Cells were primed for 24 hours prior to conducting the modified Cell Painting assay and data processing for hit identification. The CellProfiler analysis pipeline segmented nucleus and cell body accurately (**Fig. S3**). Mp-value testing (which returns a single statistic based on multidimensional Mahalanobis distance between permutations of two groups) (71) using 105 PCs to summarize 90% of the variance in the data was employed to identify priming conditions that were significantly different from the −CTL. Hierarchical clustering separated priming conditions into unique clusters (**Fig. 2A**), and we further filtered these priming conditions to identify those conditions that had all 4 replicate wells in the same cluster. Therefore, we were able to identify unique and low-variability priming conditions in an unbiased manner.

**Figure 2.**
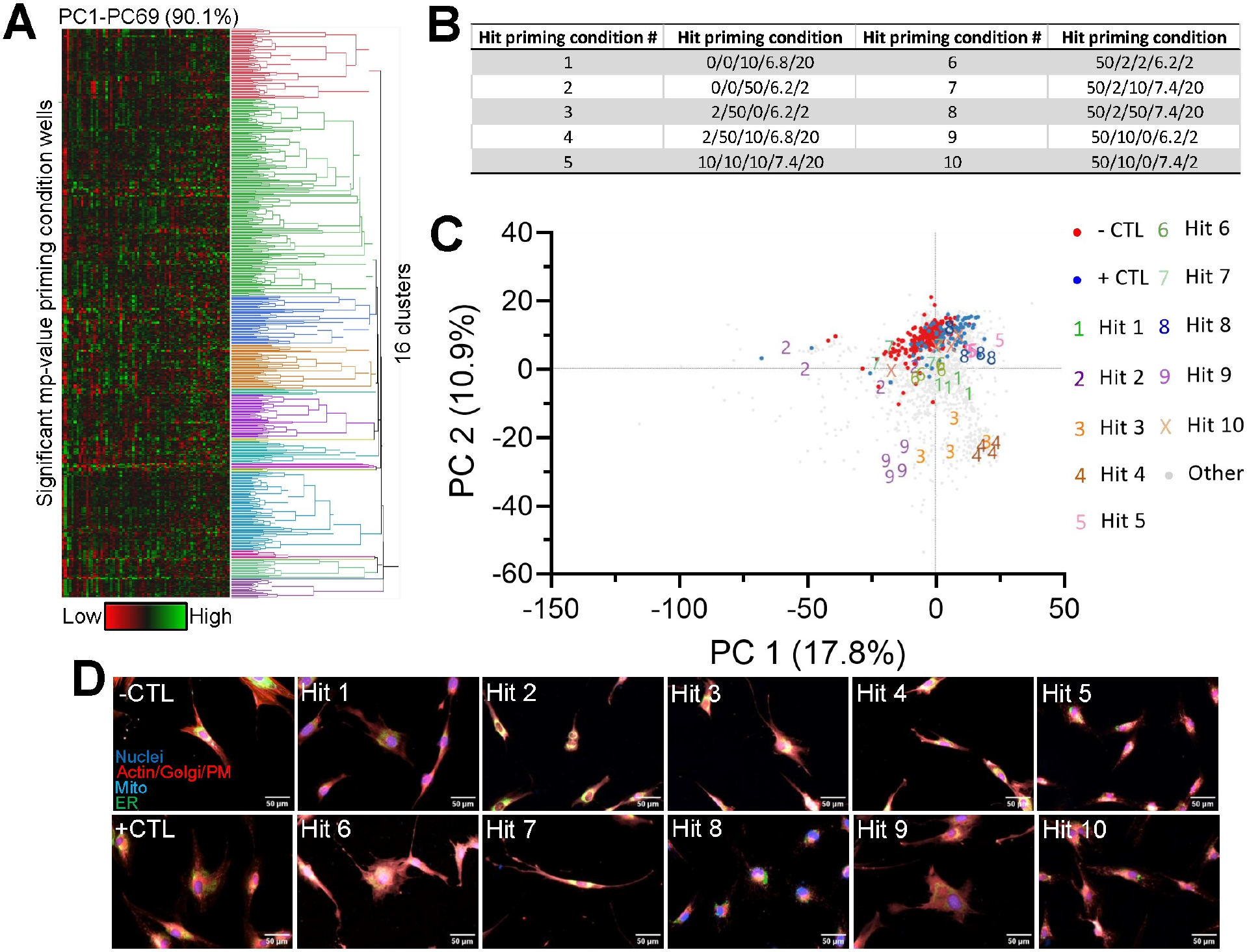
The exploratory HTS identified unique priming conditions corresponding to distinct morphological responses. **A.** Hierarchical clustering of significant mp-value wells using PCs from PCA of these wells. **B.** Exploratory HTS 10 hits by code [IFN-γ]/[TNF-α]/[IL-1β] (ng/mL)/pH/O2%. **C.** PCA (input 1,126 features) of all wells with −CTL, +CTL, and exploratory HTS 10 hits highlighted, where each point represented 1 well (1,848 total wells). **D.** Cell Painting, representative cells color composite images for −CTL, +CTL, and exploratory HTS 10 hits.

10 hits were identified in the exploratory HTS, spanning the range of cytokines and concentrations screened and roughly evenly divided between the 3 pH and 2 oxygen levels. This diversity of hits across all levels of the full factorial screen design gave confidence that the screening method identified unique manufacturing conditions (**Fig. 2B**). Replotting the data using only PC1 and PC2 (28.7% of total variance), the −CTL and +CTL groups formed distinct clusters while hits spread across PC1 and PC2 (**Fig. 2C**). Although PC1 and PC2 summarized <30% of the variance in the data (and 105 PCs were required to summarize 90% of the variance in the data) (**Fig. S4**), this is not unexpected considering the high dimensionality of the screen (1,126 features contributing to PCA). Exploratory HTS plate heatmaps colored by PC1 and cell count did not reveal patterns in values or hits of plate effects or plate layout effects (**Fig. S5**). This visual HTS quality assessment supported that hits were identified based on replicable biological phenomena as opposed to experimental artifacts. Visual assessment of the CTLs and hits confirmed qualitatively that the hit identification process is effective in identifying priming conditions corresponding to unique morphologies (**Fig. 2D**).

### Validation HTS to further refine priming hits

Next, validation HTS was performed to effectively narrow the exploratory HTS 10 hits to the 5 most unique hits and validate exploratory screen findings. Validation screening was performed identically to exploratory screening with the addition of two cell-lines: BMMSC RB71 and ADMSC RB98 were selected to test relatively low and high, respectively, indoleamine-2,3-dioxygenase (IDO) activity and T cell suppression cell-lines. The inclusion of low and high potency cell-lines attempts to address MSC functional heterogeneity and whether the hits produce universal morphological responses or if cell-line differences exist. PCA by cell-line showed the 10 hit priming conditions induced distinct morphological responses in all 3 cell-lines (**Fig. 3A**). Fewer PCs (26) were required to summarize 90% of the variance in the data in the validation HTS compared to the exploratory HTS (**Fig. S6**). Visual inspection of PCA plots, as well as desire to represent a diversity of tested conditions, informed selection of the 5 most unique hit priming conditions. Qualitative analysis of representative cells for the CTLs and hits in each cell-line confirmed uniqueness of the hits (**Fig. 3B**). Again, visual HTS quality assessment supported that hits were identified based on replicable biological phenomena (**Fig. S7**). Because of preliminary data demonstrating EVs from BMMSC RB71 modulate microglia (83), we chose this cell-line for subsequent manufacturing. Hits in **Fig. 3A** were renumbered in **Fig. 3B** for MSC-EV manufacturing according to **Table S3**. The validation HTS confirmed the exploratory HTS findings and identified the 5 most unique morphological hits to further investigate in terms of their effects on MSC-EV manufacturing.

**Figure 3.**
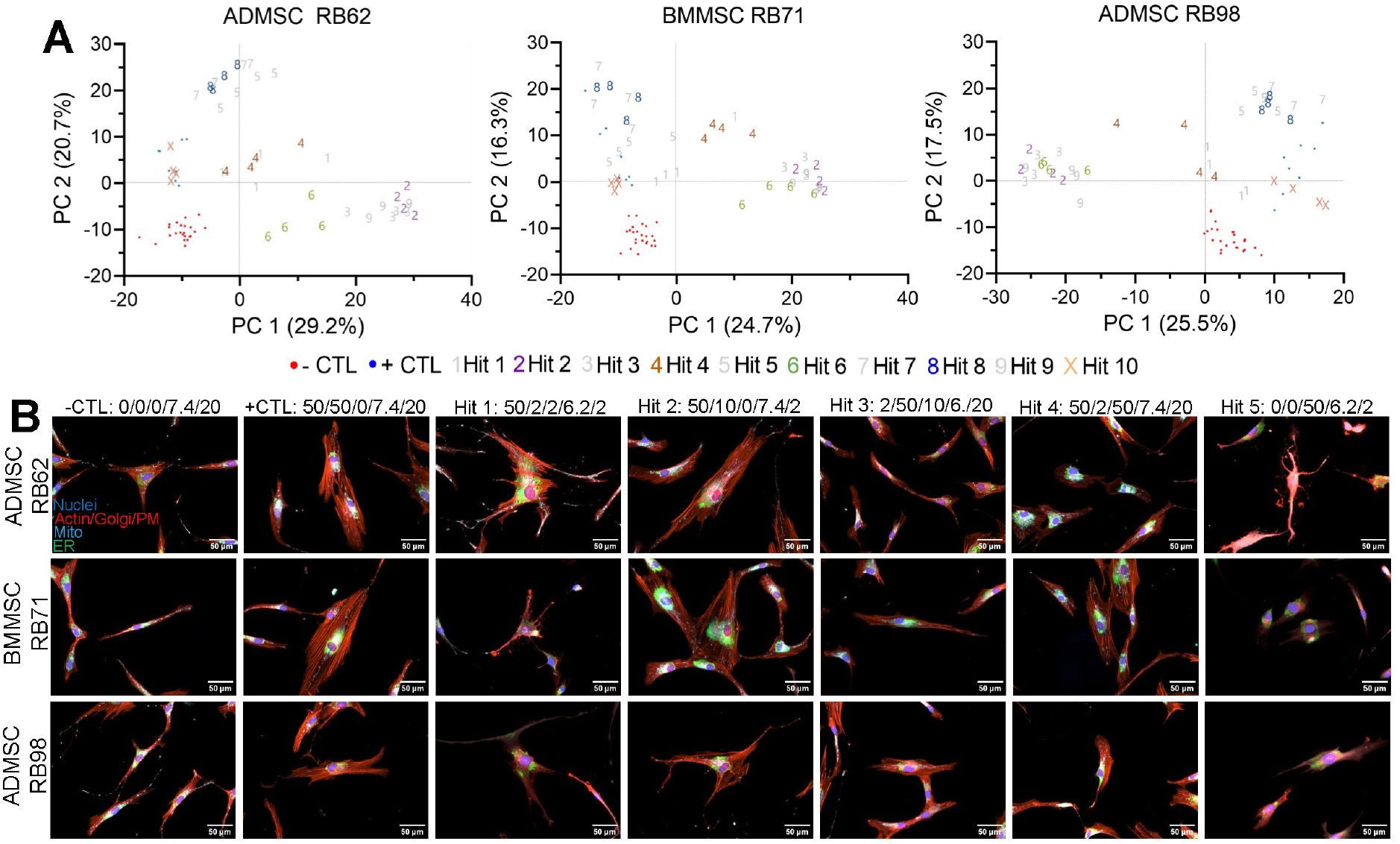
Validation HTS to further refine priming hits. **A.** PCA (input 667 features) by cell-line of all wells with - CTL, +CTL, and exploratory HTS 10 hits highlighted where each point represents 1 well (total 72 wells/cell line). **B.** Representative cells from each cell-line for −CTL, +CTL, and validation HTS 5 hits labeled by code [IFN-γ]/[TNF-α]/[IL-1β] (ng/mL)/pH/O2%. Validation HTS 5 hits (subset of 10 exploratory HTS hits) renumbered in **B** according to **Table S3**.

### +CTL, Hit 2, and Hit 4 priming conditions significantly impacted MSC-EV protein and particle yield

The 5 hit priming conditions identified through the exploratory and validation morphological screens were tested in parallel flask and bioreactor MSC-EV manufacturing, followed by evaluation of MSC-EV preparation quality (identity, purity, and potency) (**Table S2**). 0.5 L vertical wheel bioreactors were used due to their intermediate scale, plug and play implementation, and previous characterization (81). Bioreactors were shown to scale-up MSC manufacturing in a consistent manner (**Fig. S8**). Furthermore, regardless of priming condition, bioreactors qualitatively progressed similarly (in terms of microcarrier aggregation) throughout expansion and priming phases. Consistency in manufacturing was important to demonstrate as MSC-EVs were manufactured in 2 batches and including −CTL groups in each batch would allow for batch correction when comparing potency of all MSC-EV groups.

Protein yield increased significantly with priming (compared to −CTL1 within a scale) for all priming conditions in flasks (**Fig. 4A**). For bioreactor groups, protein levels were similar for +CTL and Hits 1, 3, and 5 while Hits 2 and 4 showed much higher protein levels. There were no significant differences in protein levels between flask and bioreactor groups except for Hit 4, which had significantly lower protein levels in the bioreactor compared to its respective flask. In terms of particle count, there were significant increases in both flask and bioreactor groups for +CTL, Hit 2, and Hit 4 (**Fig. 4B**). The only differences observed in particle count based on scale were for Hits 2 and 4 where bioreactor groups had slightly lower particle counts than their respective flasks. Particle diameter was significantly smaller than −CTL in the flask groups +CTL and Hits 2, 3, and 4 (**Fig. 4C**). Bioreactor groups +CTL and Hit 4 were significantly different from their respective flasks. Example distributions of particle size show the expected power-law behavior up to the lower limit of detection (75 nm) for all groups (**Fig. S9**) (82). Batch 1 and batch 2 controls (-CTL1 and −CTL2, respectively) were not statistically different in both flasks and bioreactors, further demonstrating batch-batch consistency in MSC-EV manufacturing. Pooling flask and bioreactor groups, flasks had higher average purity, although not significantly so (**Fig. 4D**). Bead-based flow cytometry was used to assess the presence of intact vesicles and a canonical EV surface marker, the tetraspanin CD81. The combination of CFSE+ staining on CD81 capture beads enabled confirmation of MSC-EV identity in terms of phenotype and intact lipid membrane. MSC-EV preparations from all groups showed CD81+ intact vesicles (**Fig. 4E**). Overall, the 5 hit priming conditions showed uniqueness in MSC-EV preparation phenotype, including with respect to scale. Notably, +CTL, Hit 2, and Hit 4 priming conditions significantly impacted MSC-EV protein and particle yield.

**Figure 4.**
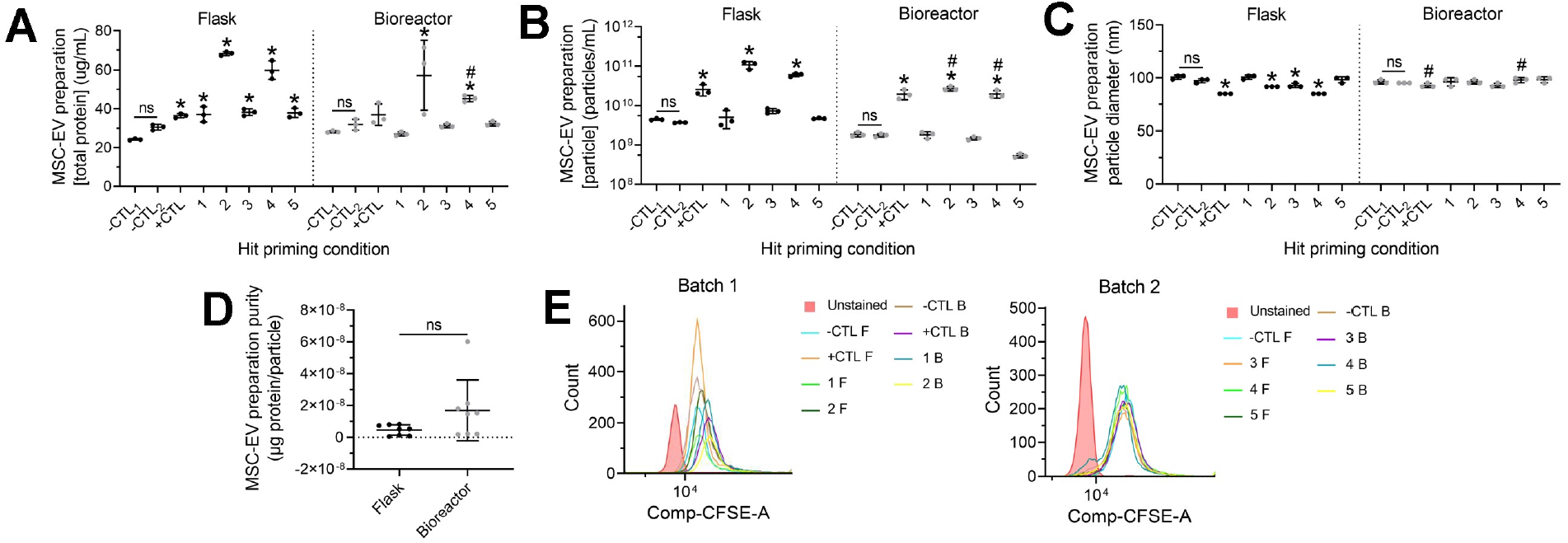
+CTL, Hit 2, and Hit 4 priming conditions significantly impacted MSC-EV protein and particle yield. **A.** MSC-EV preparation protein yield where each point represents 1 well (n = 3). **B.** MSC-EV preparation particle yield. **C.** MSC-EV preparation median particle diameter. **B-C.** Each point represents 1 Spectradyne nCS1 cartridge (n=3). **D.** MSC-EV preparation purity calculated as protein yield/particle yield (n=8). **A-D.** Graphed as mean and standard deviation. **A-C.** *p<0.05 versus −CTL1 within a scale, 2-way ANOVA with Dunnett’s multiple comparisons test, #p<0.05 versus respective flask group, 2-way ANOVA with Šidak’s multiple comparisons test. **D.** ns, paired t-test. **E.** MSC-EV bead-based flow cytometry for batch 1 and batch 2 MSC-EV preparations. Abbreviations: F: flask; B: bioreactor.

### Priming conditions in a bioreactor resulted in differential MSC-EV microglia modulation

Potency of MSC-EV preparations was then tested in a novel image-based microglia modulation assay in which MSC-EV preparations were administered concurrently with inflammatory cytokines IFN-γ and TNF-α to a human microglia cell-line (C20) and shifts in microglia morphology were quantified using high dimensional morphological analysis. Microglia morphology, represented by composite morphological score PC1 comprising 21 morphological features, was used as an output of MSC-EV potency (83) as changes in microglia morphology occur when they are aberrantly activated in neurodegenerative diseases (84). *In vitro*, microglia become more elongated in response to inflammatory signals present in neuroinflammation, such as TNF-α (85–87). In this assay, higher PC1 microglia morphology scores are associated with stimulated, activated microglia morphology (i.e., ‘inflammatory’) while lower PC1 scores are indicative of unstimulated, inactivated microglia (i.e., ‘resting’). Thus, lower PC1 scores are indicative of more ‘potent’ MSC-EVs. All treatments were significantly lower than the stimulated microglia CTL (*p<0.05, **Fig. 5B**) as the overall morphology shifted towards the unstimulated microglia CTL. Although significantly different from the stimulated microglia CTL, MSC-EVs from bioreactor groups −CTL1, −CTL2, and Hits 3 and 5 (#p<0.05) did not have as significant of an effect on microglia morphology as respective flask groups (i.e., were less potent). The most potent hit priming conditions in the bioreactor (being the only conditions performing similarly to the +CTL and respective flask groups) were Hits 2 and 4 as they possessed the lowest PC1 scores while Hits 1, 3, and 5 were less potent as they possessed significantly higher PC1 scores ($p<0.05). Visual assessment of images confirmed microglia increase in size and elongation with stimulation, and EV treatment reverses this trend towards the unstimulated microglia control. (**Fig. 5A, C**). In our MSC-EV potency assay, the bioreactor groups remained most relevant for further investigation due to the requirement of increased scale for clinical translation. Furthermore, the assay served as a functional screen of priming conditions, necessitating validation of these priming conditions. In summary, morphological screening identified priming conditions in a bioreactor that resulted in differential microglia modulation with +CTL, Hit 2, and Hit 4 resulting in the greatest shift (although Hit 2 and Hit 4 were not significantly different from +CTL) in microglia morphology towards the inactivated morphology (i.e., lowest PC1 scores in the bioreactor group).

**Figure 5.**
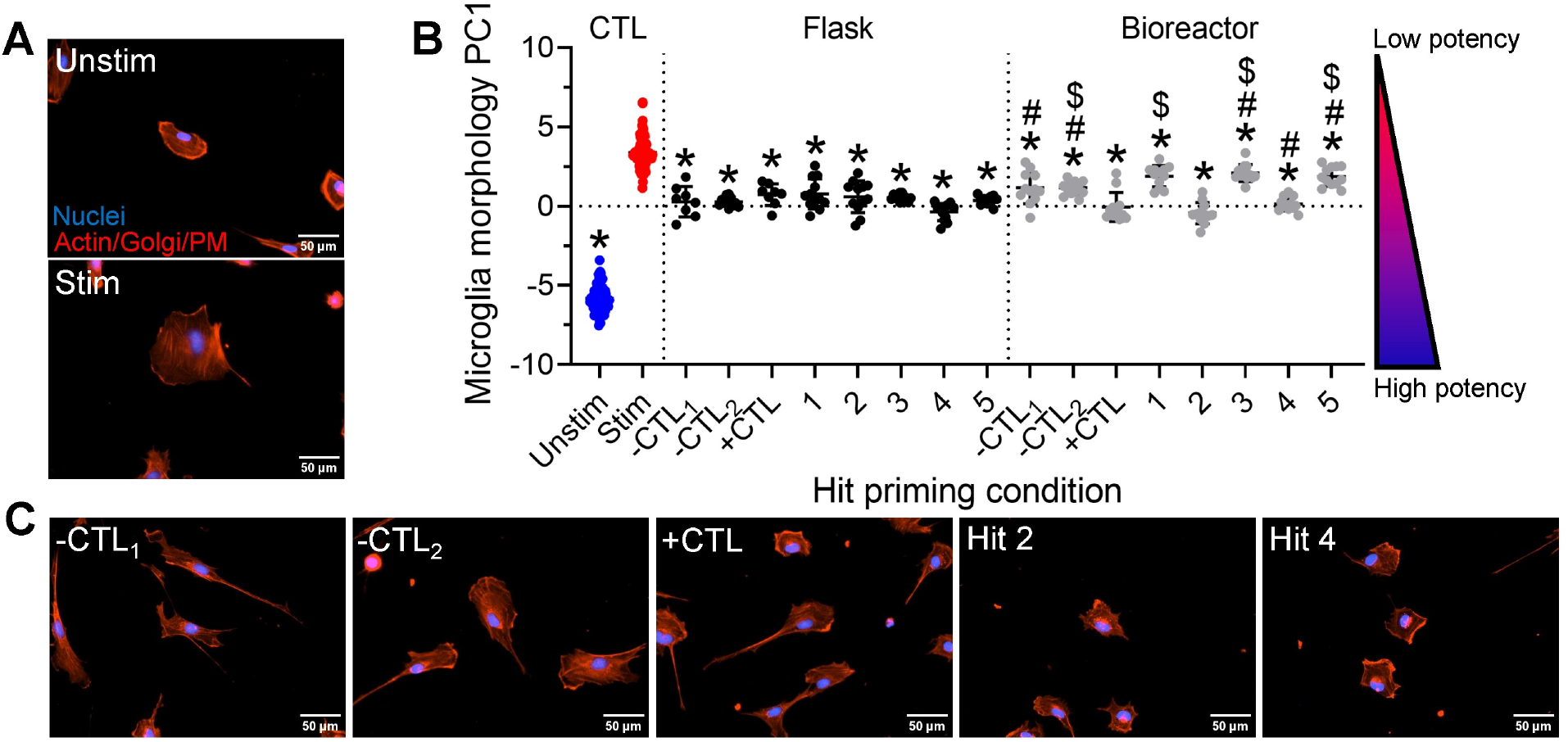
Priming conditions in a bioreactor resulted in differential MSC-EV microglia modulation. **A.** Unstim (microglia unstimulated CTL) and Stim (microglia stimulated CTL) representative cells color composite images. **B.** Morphology-based microglia modulation assay. Lower microglia morphology PC1 scores indicate more potent MSCEVs. Graphed as mean and standard deviation where each point represents 1 well, n = 72 (unstim, stim), n = 8-12 (all other groups), *p<0.05 vs stim, $p<0.05 vs +CTL within same scale, Brown-Forsythe and Welch ANOVA with Dunnett’s T3 multiple comparisons test, #p<0.05 vs respective flask group, ordinary 2-way ANOVA with Šidak’s multiple comparisons test. **C.** MSC-EV treatment representative cells color composite images.

### Validation MSC-EV manufacturing confirmed priming hits significantly modulate microglia

Although no bioreactor groups had enhanced potency (lower PC1 scores) compared to respective flask groups, we chose to follow-up our initial MSC-EV manufacturing study with a focused bioreactor only study due to the following considerations: 1) our bioreactor MSC-EV preparations with the highest potency (+CTL, Hit 2, and Hit 4) were similar to flask groups for the same priming conditions and 2) increased scale for clinical translation necessitates bioreactor manufacturing systems. A validation MSC-EV manufacturing was performed to confirm high potency hit priming conditions, as well as explore whether changing bioreactor parameters such as microcarrier (µC) density impact MSC-EV quality. Microcarrier concentration was increased by a factor of 15.625X to ensure bioreactors were operating at capacity in terms of cell expansion and MSC-EV yield and to assess hit performance across changes in an important bioreactor parameter. For µC_low_, there was significantly higher protein yield in Hits 2 and 4 compared to the +CTL (*p<0.05, **Fig. 6A**). Hits 2 and 4 were also significantly different in µC_high_, although Hit 4 was lower than the +CTL here. Only the +CTL and Hit 2 were significantly different from their respective µC_low_, both being higher (#p<0.05). Particle count showed similar trends in both µC_low_ and µC_high_ (**Fig. 6B**). However, there were no significant differences between the hits and the +CTL in either case and (despite relatively high variability) the +CTL was the only in µC_high_ that was significantly higher than its respective µC_low_, although µC_high_ were generally higher than µC_low_. There were no significant differences between the +CTL and any hits for particle diameter, although +CTL µC_high_ was significantly different from its respective µC_low_ (**Fig. 6C**). In terms of purity, although greater variability existed for µC_low_, these groups were not significantly different from µC_high_ (**Fig. 6D** and **Fig S10**).

**Figure 6.**
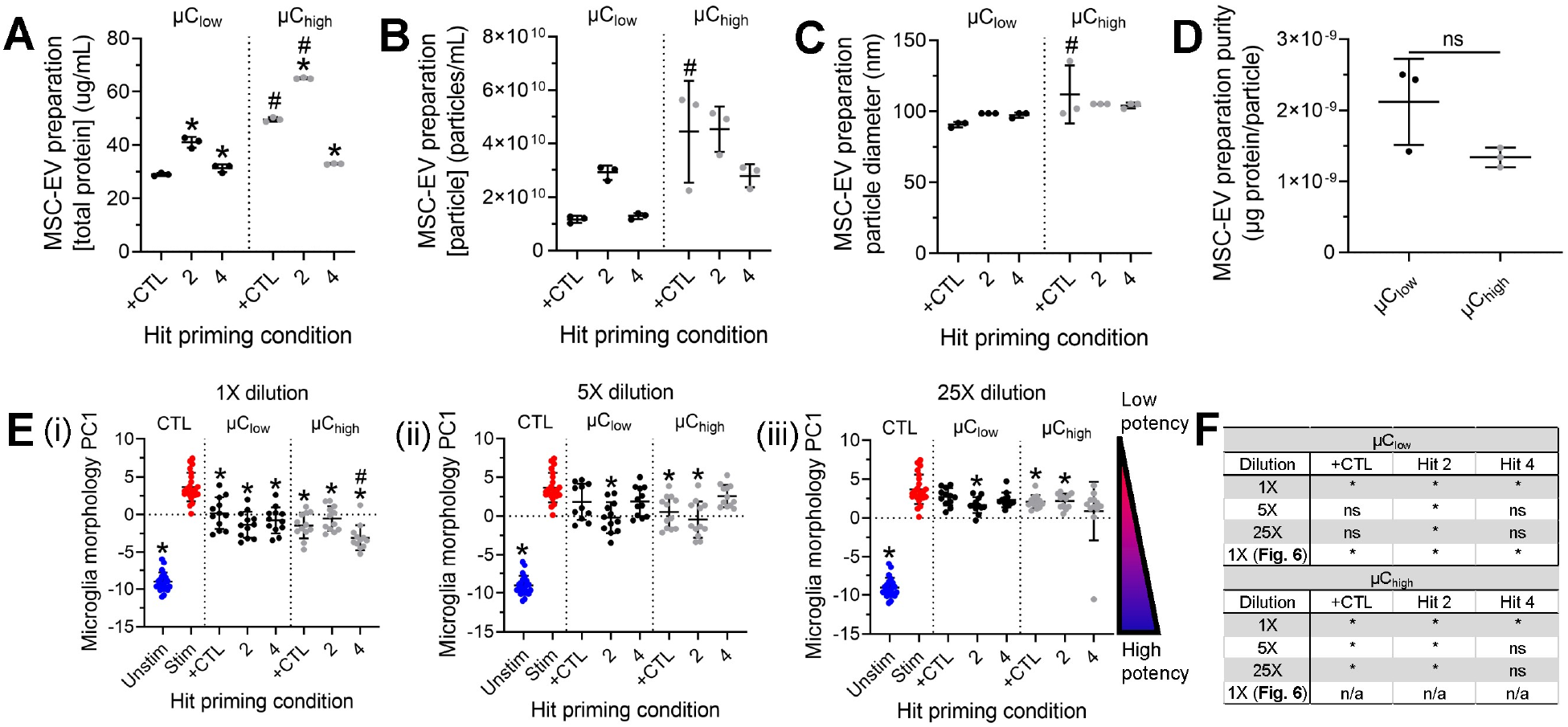
Validation MSC-EV manufacturing confirmed priming hits significantly modulate microglia. **A.** MSCEV preparation protein yield where each point represents 1 well (n = 3). **B.** MSC-EV preparation particle yield. **C.** MSCEV preparation median particle diameter. **B-C.** Each point represents 1 Spectradyne nCS1 cartridge (n=3). **D.** MSCEV preparation purity calculated as protein yield/particle yield (n=3). **A-D.** Graphed as mean and standard deviation. **A-C.** *p<0.05 versus +CTL within μC, 2-way ANOVA with Dunnett’s multiple comparisons test, #p<0.05 versus respective μC, 2-way ANOVA with Šídák’s multiple comparisons test. **D.** ns, paired t-test. **Ei-iii.** Morphology-based microglia modulation assay with 3 MSC-EV preparation dilutions (1X, 5X, and 25X). Lower microglia morphology PC1 scores indicate more potent MSC-EVs. Graphed as mean and standard deviation where each point represents 1 well, n = 23 (unstim), n = 24 (stim), n = 11-12 (all other groups), *p<0.05 vs stim, ns vs +CTL within μC, Brown-Forsythe and Welch ANOVA with Dunnett’s T3 multiple comparisons test, #p<0.05 vs respective μClow, ordinary 2-way ANOVA with Šídák’s multiple comparisons test. **F.** Table summarizing statistical results from **Fig. 5B** and **Fig. 6Ei-iii**.

The microglia morphology potency assay was performed (as in **Fig. 5**) with the addition of multiple MSC-EV dilutions (1X, 5X, and 25X) as a means to assess MSC-EV dose response (**Fig. 6E**). At 1X, all treatments performed significantly better than the stimulated microglia CTL (*p<0.05) and only Hit 4 in µC_high_, possessing the highest potency overall (lowest PC1 score), was more potent than its respective µC_low_ (#p<0.05). No groups were significantly different from the +CTL. However, at the 5X dilution, only Hit 2 in both µC groups and the +CTL in µC_high_ were significantly different from the stimulated microglia CTL. Furthermore, this trend was observed at 25X dilution. The table in **Fig. 6F** summarizes the statistical results in **Fig. 5B** and **Fig. 6E** and highlights that Hit 2 was significantly different from the stimulated microglia CTL across all dilutions and µC groups, unlike the +CTL and Hit 4, indicating a potentially more cost effective and robust priming condition. Overall, µC did not have a consistent impact on MSC-EV preparation quality. The consistently high potency across multiple experiments, dilutions, and µC suggests Hit 2 as an ideal priming condition for bioreactor-based manufacturing of MSC-EVs for microglia modulation.

### Priming condition and µC impacted MSC-EV lipid composition

Finally, lipidomics was performed using liquid chromatography-mass spectrometry (LC-MS) on MSC-EV preparations from the validation manufacturing to gain insight into metabolic regulation of MSC-EV production in response to priming as well as mechanism of action in terms of modulating microglia. Both positive and negative electrospray ionization were used for data collection, with data analysis resulting in annotation of 62 individual lipid species in 8 lipid classes: bis(monoacylglycero)phosphate (BMP), phosphatidylethanolamine (PE), phosphatidylglycerol (PG), vinyl ether-linked (plasmalogen) phosphatidylethanolamine (PE P), diacylglycerol (DG), phosphatidylcholine (PC), alkyl ether-linked (plasmanyl) phosphatidylcholine (PC O), and sphingomyelin (SM). Retention time and MS/MS spectra were used to confirm lipid identities (**Fig. S11-S15**). The predominant trends included increased lipid intensity with increased µC for at least one priming condition in every lipid species, many times for two or all conditions. Hit 2 showed significantly higher lipid intensity than +CTL for 47 species in µC_low_ and 37 species in µC_high_. Hit 4 showed significantly different lipid intensity from +CTL for only three lipid species in µC_low_ and significantly lower lipid intensity than +CTL for all 62 lipid species in µC_high_ (**Fig. 7A-B**). Thus, Hit 2 demonstrated ubiquitous significant increases in lipid intensity compared to +CTL. Analysis of the lipidomic data using PCA enabled assessment of the overall lipid profiles with PC1 and PC2 separating conditions based on µC and priming condition (**Fig. 7C**). PC1 accounted for 90% of the variance in the data and we therefore performed statistical comparisons of the groups using this metric as a composite lipidomic score. All µC_high_ were significantly different from their respective µC_low_, and Hit 2 was different from +CTL in both µC groups, whereas Hit 4 was only different from the +CTL in µC_high_, evidencing a potential interaction between priming condition and µC. Looking only within µC_high_ showed a similar result (**Fig. 7D**). Thus, Hit 2 had a distinct lipidomic profile, corroborating its uniqueness in the potency assay.

**Figure 7.**
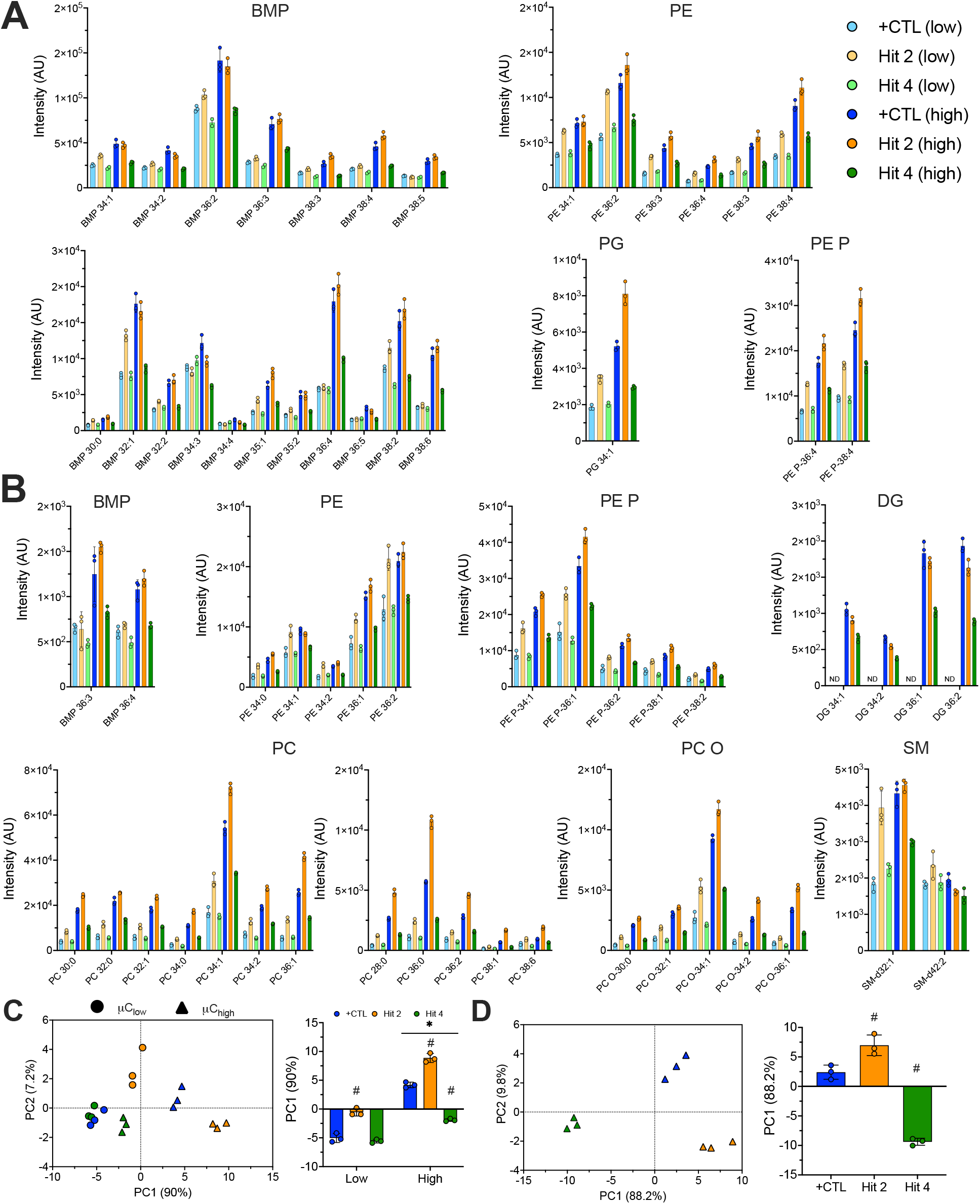
Priming condition and µC impact MSC-EV lipid content. **A.** MS negative ionization mode lipid intensities. **B.** MS positive ionization mode lipid intensities. **A-B.** #p<0.05 vs respective µClow, $p<0.05 vs +CTL within a µC, ordinary 2-way ANOVA with Dunnett’s multiple comparisons test. **C.** PCA on all positive mode lipid species intensities (excluding DG lipid species because they were below the limit of detection in µClow), *p<0.05 vs µClow within a priming condition, #p<0.05 vs +CTL within µC, Brown-Forsythe and Welch ANOVA tests with Dunnett’s T3 multiple comparisons test. **D.** PCA on all negative and positive mode lipid intensities for µChigh, #p<0.05 vs +CTL. **A-D.** Graphed as mean and standard deviation where each point represents 1 technical replicate (n=3, except for Hit 4 (low) positive mode, n=2).

### Correlating MSC-EV preparation quality and potency

Because there is no normalization of MSC-EV preparations in the potency assay (i.e., no adjustment of MSC-EV preparations to equalize dose in terms of particle count or protein mass), performance of different MSC-EV preparations can be directly interpreted considering our quality metrics. We combined data from all 22 MSC-EV preparations to explore these relationships. Protein and particle concentration were significantly correlated (**Fig. 8A**), although neither were predictive of potency measured by PC1 (**Fig. 8Bi-ii**). Notably, the logarithm of particle concentration strongly correlated with potency (**Fig. 8Ci**), but the logarithm of protein concentration did not (**Fig. 8Cii**).

**Figure 8.**
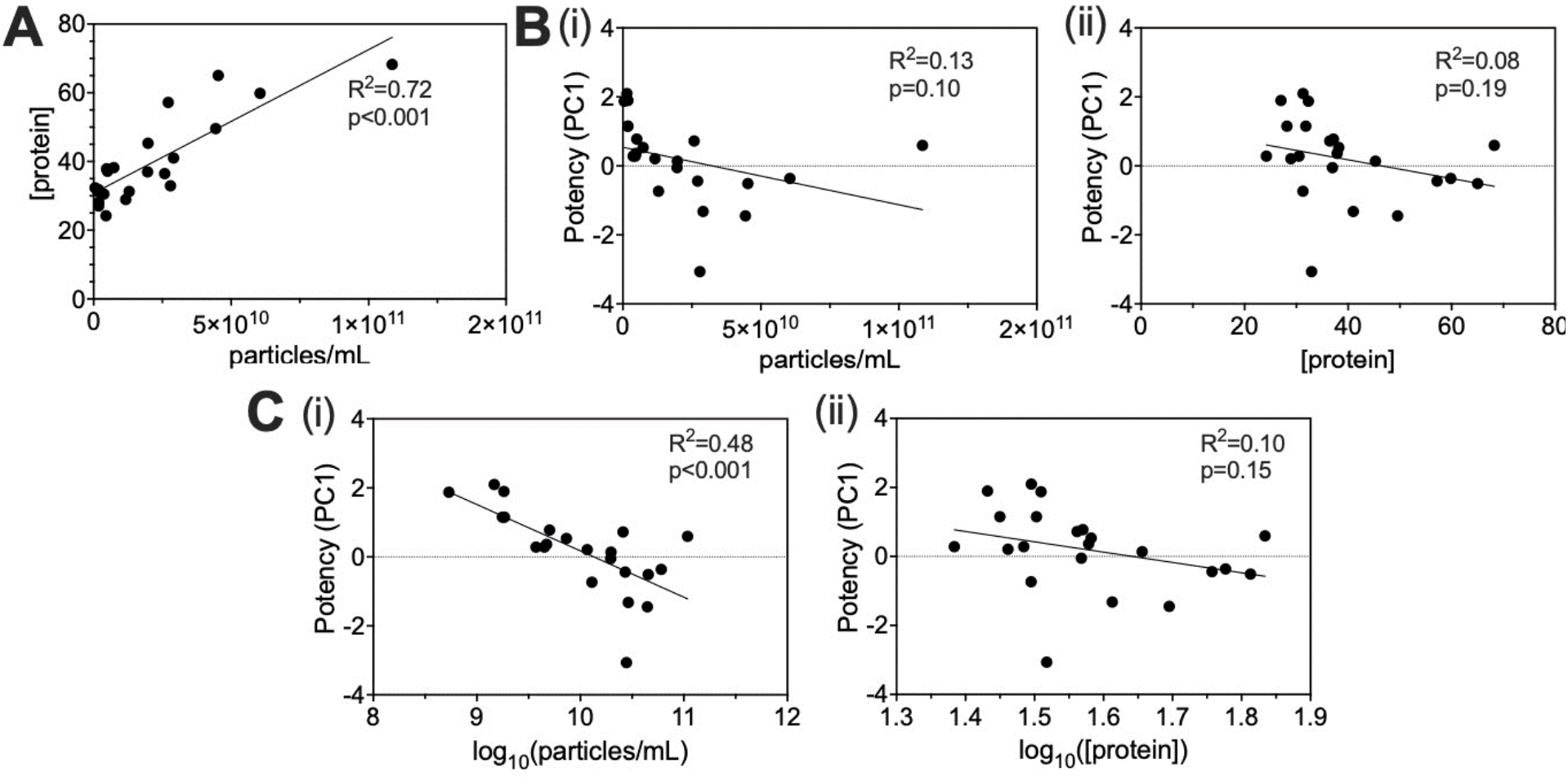
Correlating MSC-EV preparation quality and potency. **A.** Protein yield-particle yield linear regression. **Bi.** Microglia morphology PC1 score-particle yield linear regression. **Bii.** Microglia morphology PC1 score-protein yield linear regression. **Ci.** Microglia morphology PC1 score-log10(particle yield) linear regression. **Cii.** Microglia morphology PC1 score-log10(protein yield) linear regression. **A-C.** Combined quality metrics from 22 MSC-EV preparations.

## DISCUSSION

MSC priming is an emerging manufacturing strategy that has the potential to address challenges of functional heterogeneity and translate promising MSC-based therapies to the clinic. However, priming strategies have not been comprehensively explored, including in their interaction with clinically relevant manufacturing formats. We demonstrated morphological screening as a tool to inform and enhance production of immunomodulatory MSC-EVs in a bioreactor. After validating novel, potent priming conditions for microglia modulation, we probed mechanisms of the MSC functional response to priming and mechanisms of MSC-EV microglia modulation via MSC-EV lipidomics.

To our knowledge, this is the most comprehensive screen of MSC priming conditions. Most previous studies have investigated the combination of only two signals, potentially missing more potent or cost-effective conditions (26, 27, 31, 35–38, 40, 51, 88). Furthermore, this work is a pioneering application of Cell Painting to screen cell therapy response to changes in manufacturing conditions and one of few studies leveraging morphology as a screening tool to specifically assess MSC response to changes in manufacturing conditions (26, 61, 62, 89–94). Although Cell Painting involves staining of intracellular structures rather than specific biomarkers, measured changes in morphology have been broadly related to function in various cell types: neural cell differentiation potential in neural stem cells (95) and metastatic potential in cancer cells (96, 97), as well as MSC immunomodulatory function (26, 61, 62). In this study, ADMSC RB62 consistently increased in size (cell area) and decreased in elongation (cell aspect ratio) and complexity (cell form factor) with priming. Increased cell area with priming agrees with previous studies using 4 MSC donors (26) and 6 MSC donors (61); however, the decreased elongation and complexity we observed differs from previous trends, showing that although functional heterogeneity decreases in response to priming, cell-line dependent morphological heterogeneity still exists. Furthermore, structures stained by Cell Painting directly impact EV biogenesis, so changes in morphology can be mechanistically related to MSC response to priming. For instance, actin and actin-associated proteins (e.g., myosins) are packaged into EVs (98) and actin may play a role in formation and trafficking of EVs, microvesicles, and multivesicular bodies (99–103). Also, plasma membrane protrusions have been shown to be specific platforms for EV shedding that require actin rearrangement (100–102, 104–106). Thus, larger-scale changes in actin structure that we measure may be indicative of changes in MSC-EV production.

Importantly, we investigated the effects of larger-scale manufacturing on MSC functional response to priming and found that hit priming conditions differentially impacted MSC-EV identity and potency in bioreactors. This work represents one of few studies investigating MSC priming in a bioreactor (107) and likely the first study evaluating EVs in this context. Our result that +CTL, Hit 2, and Hit 4 significantly impacted MSC-EV protein and particle yield agrees with previous evidence of increased MSC-EV protein and particle yield with priming as well as a synergistic effect of combinatorial priming (27, 29, 31, 37, 44–47, 49–52, 108–112). Combined hypoxia and cytokine priming has been shown to significantly improve MSC immunomodulatory function compared to either signal alone (31, 108). Our Hit 2 shows a similar phenomenon in the EVs of hypoxia and cytokine primed MSCs. Both hypoxia and cytokines are known to independently shift MSC metabolism to aerobic glycolysis (108, 113); metabolic reconfiguration induced by dual hypoxia and cytokine priming may fuel changes in MSC-EV production and lipid profile related to potency. The protein, particle count, and particle size we observed are within the range seen in similar studies (27, 37, 44, 46, 47, 49–53, 109–112). The lower purity in our MSC-EV preparations compared to other studies using MSCs (114, 115) may be explained by differences in cell manufacturing and MSC-EV isolation and analysis methods. Although we observed significant decreases in particle diameter for several groups, significant changes in median particle diameter do not necessarily signify significant changes in particle diameter distribution, a more meaningful quantification of changes in EV biogenesis. Stable expression of CD81 across priming conditions also agreed with the similar studies cited previously. We did not observe a consistent significant effect of scale or µC on MSC-EV protein and particle yield or potency. MSC-EVs have been manufactured in larger-scale formats such as the Nunc Cell Factory, 3D microcarrier-based systems, hollow fiber bioreactors, vertical wheel bioreactors, and stirred tank bioreactors with 2.2-20 fold increases in yield (49, 114–119). However, we expanded MSCs in flasks prior to MSC-EV manufacturing to obtain a sufficient number of cells for seeding parallel flasks and bioreactors; thus, generally similar yield in terms of µg protein/mL and particles/mL between our flasks and bioreactors may be due to the need for MSCs to adapt to the changed format of bioreactors, impacting MSC-EV production compared to flasks. For instance, it has been shown that bioreactor and 3D microcarrier culture place new demands on MSCs in terms of metabolism, mechanical forces, and interactions with substrates; all of these factors can eventually impact MSC function (120). Ultimately, inconsistencies in MSC tissue sources, bioreactor formats, priming approaches, and MSC and EV analytical methods complicate comparisons between studies and reinforce the need for standardization (121).

EVs from MSCs primed with +CTL, Hit 2, and Hit 4 significantly modulated microglia in both the initial and validation manufacturing experiments. This finding agrees with the general consensus that MSC preconditioning strategies impact MSC-EV cargo (e.g., mRNA, miRNA, cytokines, and proteins) and function. Our novel microglia modulation assay provides a neuroinflammation-relevant context for assessing MSC-EVs from different priming conditions. MSC-EV modulation of microglia has been explored both *in vitro* and *in vivo*: ADMSC-EVs significantly decreased TNF-α secretion by microglia stimulated with LPS or amyloid-beta (Aβ), a hallmark peptide of Alzheimer’s Disease (122). In a triple transgenic mouse model of Alzheimer’s Disease, IFN-γ/TNF-α primed MSC-EVs delivered intranasally reduced microglia Iba-1 and CD68 (activation markers) expression and soma size, signifying an overall suppression of microglia activation (123). Our microglia morphology assay represents a new approach for screening the effects of changes in manufacturing conditions on EV potency and may be further developed into a generalizable EV release assay (124, 125). Selecting priming conditions corresponding to unique morphologies in the exploratory and validation HTS unsurprisingly resulted in a range of potency in the microglia modulation assay. Validation of +CTL, Hit 2, and Hit 4 as high potency priming conditions demonstrated the ability of morphological screening combined with priming as an approach to reduce not only MSC functional heterogeneity, but also EV functional heterogeneity (126, 127).

After considering the primary goal of enhanced potency induced by novel priming conditions, we considered secondary goals of decreased cost and increased consistency of these select potent priming conditions. Despite no consistent significant increase in MSC-EV yield in bioreactors, principles of scale-up (as opposed to scale-out) reduce manufacturing and personnel costs per EV and simplify handling of the process by technicians (128). Furthermore, performance of a hit across multiple dilutions is appealing economically and in terms of robustness, allowing identification of Hit 2 as a singular ideal priming condition for our application.

Although there was a significant correlation between particle count and potency (on a log scale), increasing particle count of low potency MSC-EV preparations through additional concentration to increase potency would result in significantly greater manufacturing costs as additional cellular material, media, and equipment would be needed. As an example, a ‘low potency’ MSC-EV preparation with particle concentration of ∼1E9 particles/mL (upper left data points in **Fig. 8C**) would need to be concentrated nearly 100X (assuming dose follows the trendline) to become as potent as a ‘high potency’ MSC-EV preparation (lower right data points in **Fig. 8C**). Therefore, we believe considering each MSC-EV preparation as derived from the same starting cell equivalent (from the same media volume and concentrated by the same factor) enables us to effectively compare effects of priming and scale on MSC-EV quality without normalizing to particle count or protein content. Finally, because priming results in changes in MSC-EV composition (as with lipidomics), normalization to particle count or protein could introduce artifacts and complicate interpretation of the results considering MSC-EV functional heterogeneity is well-established not only between batches but also within a batch (129).

Besides significantly impacting total MSC-EV yield (protein and particle concentrations), we showed that priming condition and µC impacted MSC-EV lipid content. Lipidomics is a powerful approach to mechanistically probe MSC-EV function (130, 131), including in response to priming (132). MSC metabolism shifts to aerobic glycolysis after stimulation with inflammatory signals and this shift is necessary for observed increases in MSC immunomodulatory function *in vitro* (53). Several MSC lipidomics studies have demonstrated changes in lipid profiles with priming (133–136). For instance, BMMSCs from five donors cultured under hypoxia increased total triglycerides, fatty acids, and DGs compared to controls (133). However, fewer studies exist that investigate the lipid profile of MSC-EVs. In our study, we saw significantly increased lipid intensities (particularly for Hit 2) in many lipid species of the classes PC and PE, both major cellular membrane components (137). PC and PE content in exosomes has also been documented in mast and dendritic cells (138). Changes in these lipid classes could be related to changes in the cell membrane permitting the morphological response to priming and increased EV release. In fact, lipidomics offers mechanistic insight into morphological changes in response to priming. Sphingosine-1-phosphate (S1P), a catabolite of ceramide, is a well-documented regulator of cytoskeletal reorganization in a cell type-specific manner (139, 140). Both S1P and ceramide are intermediates in sphingolipid metabolism, along with sphingomyelin (141), which we observed in MSC-EVs. Enrichment of lipids in EVs (including SMs, sphingolipids, and ceramides) compared to their originating cells has been observed in many cell types, including human prostate carcinoma cells, immortalized mouse oligodendroglial cells, and B cells (142–144).

To our knowledge, this was the first study demonstrating the presence of BMP lipids in MSC-EVs. BMP is thought to be a lipid biomarker specific to EVs from the endosomal source (regulated by the endosomal sorting complexes required for transport, or ESCRTs (145)) (146, 147), as it plays a role in intraluminal vesicle biogenesis and is not found in the plasma membrane, meaning it is not present in microvesicles (148, 149). Our unique identification of BMPs in MSC-EV preparations is consistent with the fact that we processed conditioned media using a 0.2 µm filter prior to UC, thus enriching for MSC-EVs in the size range of exosomes (30-150 nm (150)), which uniquely contain BMPs. BMPs are known to be present at higher levels in macrophages and microglia due to their role in endolysosomal integrity and function (151). Changes in autophagy have been implicated in neurodegenerative diseases, including Alzheimer’s Disease, resulting in accumulation of products typically degraded (e.g., protein aggregates such as Aβ) (152, 153). In neuronal cells, endolysosomal stress induced by phosphatidylinositol 3-phosphate (PI3P) deficiency promoted the secretion of exosomes enriched in BMP and undigested lysosomal substrates, including amyloid precursor protein C-terminal fragments (154). Although it remains unknown if BMP is a mediator or marker for diseases featuring endolysosomal dysfunction, significantly higher BMP levels in Hit 2 MSC-EVs may suggest a role of transferred BMP in reverting activated microglia back to the inactivated state, possibly through intervention at endolysosomes.

Our EV lipidomics results can also offer mechanistic insight into MSC-EV function (including in response to priming) on target cells, including microglia. To our knowledge, the only other study investigating MSC-EV lipidomic response to priming stimulated BMMSCs with 1% [O2] and serum deprivation and found enrichment of PE in exosomes (132), a class we also observed to be not only present, but significantly elevated in Hit 2. However, MSC-EVs have demonstrated remodeling of lipid metabolism in other cell types: treatment of human regulatory macrophages with BMMSC-EVs resulted in significantly reduced secretion of the pro-inflammatory cytokines IL-22 and IL-23 and secretion of PGE2, which the authors interpreted as a sign of lipid mediator class switching from inflammatory lipid mediators to specialized pro-resolving lipid mediators (SPMs), an example of lipid regulation of MSC-EV immunomodulatory function (155). Concerning microglia, one study alluded to the potential therapeutic effects of MSC-EV transfer of PCs on neuroinflammation by demonstrating *in vitro* treatment of LPS stimulated murine primary microglia with PC(16:0/16:0) significantly decreased IL-1β secretion (156). Similarly, exogenous supplementation of docosahexaenoic acid (DHA), an n-3 polyunsaturated fatty acid (n-3 PUFA), targeted LPS receptor surface location, reducing LPS induced IL-1β and TNF-α secretion in murine microglia BV2 (157). Maresin 1, a SPM derived from DHA, increased human microglial cell-line CHME3 phagocytosis of Aβ42 and decreased expression of the pro-inflammatory markers CD86, CD11b, and MHC-II when added to unstimulated microglia *in vitro* (158). Taken together, exogenous fatty acids and their derivatives can regulate microglia function, possibly via promoting healthy mitochondria bioenergetics which favor improved phagocytosis, generally considered beneficial in neuroinflammatory diseases such as Alzherimer’s Disease (159). Membrane phospholipids (which we observed significant levels of in Hit 2) are sources of DHA, arachidonic acid, and eicosapentaenoic acid, which through diverse enzymatic activity can produce the SPMs lipoxins, resolvins, protectins, and maresins that importantly regulate inflammatory response (160–163). These studies of lipid regulators of microglia inflammatory phenotype show MSC-EV lipidomics may grant valuable insight into MSC-EV metabolic mechanisms of action in neuroinflammatory contexts.

## CONCLUSION

We established a novel, exploratory application of Cell Painting in the most comprehensive screen of MSC priming conditions and are one of few groups to assess the effects of priming in bioreactors. Our application of a more commonly employed high throughput drug screening approach to a cell manufacturing context enabled identification of unique priming conditions that corresponded to a range of potency of MSC-EVs manufactured in bioreactors. This approach is generalizable to different aspects of cell manufacturing and can be readily adapted to different therapeutic cell-types and manufacturing reagents/culture platforms (e.g., media, biomaterials). Establishing a baseline response of MSCs to priming in bioreactors will enable further refinement of manufacturing methods based on observed differences in MSC-EV production, modulation of microglia, and lipidomics. Additional -omics studies (e.g., transcriptomics, proteomics) in combination with pathway analysis will contribute to improved MSC-EV CQAs and understanding of MSC-EV mechanisms of action in models of immune disease that will further inform approaches to enhance MSC-EV function.

## Abbreviations

ADMSC: adipose-derived mesenchymal stromal cell
Aβ: amyloid-beta
BMMSC: bone marrow-derived mesenchymal stromal cell
BMP: bis(monoacylglycero)phosphate
CQA: critical quality attribute
DG: diacylglycerol
DHA: docosahexaenoic acid
DOE: design of experiments
ESCRT: endosomal sorting complexes required for transport
EV: extracellular vesicle
GM: growth media
HCI: high content imaging
HPC: high performance computing
HTS: high throughput screen
IDO: indoleamine-2,3-dioxygenase
IFN-γ: interferon-gamma
IL: interleukin
ISCT: International Society of Cell and Gene Therapy
mp-value testing: multidimensional perturbation value testing
MS: mass spectrometry
MSC: mesenchymal stromal cell
PC: principal component
PC: phosphatidylcholine
PC O: alkyl ether-linked (plasmanyl) phosphatidylcholine
PCA: principal component analysis
PE: phosphatidylethanolamine
PE P: vinyl ether-linked (plasmalogen) phosphatidylethanolamine
PG: phosphatidylglycerol
PGE2: prostaglandin E2
PI3P: phosphatidylinositol 3-phosphate
PUFA: polyunsaturated fatty acid
S1P: sphingosine-1-phosphate
SM: sphingomyelin
SPM: specialized pro-resolving lipid mediator
TNF-α: tumor necrosis factor-alpha
UC: ultracentrifugation

## CRediT Authorship Contribution Statement

AML: Conceptualization, Data Curation, Formal Analysis, Investigation, Methodology, Visualization, Writing – original draft, Writing – review & editing; TMS: Data curation, Formal analysis, Investigation, Methodology; KRD: Data curation, Formal analysis, Methodology, Writing – review & editing; MGM: Data curation, Investigation, Methodology; HMH: Data curation, Formal analysis, Investigation, Methodology, Writing – review & editing; JMC: Data curation, Formal analysis, Investigation, Methodology, Writing – review & editing; KMH: Conceptualization, Formal analysis, Project administration, Resources, Supervision, Writing – review & editing; RAM: Conceptualization, Data curation, Formal analysis, Funding Acquisition, Investigation, Project Administration, Resources, Supervision, Writing – review & editing.

## Acknowledgements

Drs. Ty Maughon and Seth Andrews helped with the expansion and banking of all MSC lines used in the study. We consulted with Drs. Anne Carpenter, Beth Cimini, Shantanu Singh (Broad Institute) in the planning and development of our HTS approach. This study was supported in part by resources and technical expertise from the Georgia Advanced Computing Resource Center, a partnership between the University of Georgia’s Office of the Vice President for Research and Office of the Vice President for Information Technology. We consulted with Akash Ramachandran, Ravi Jyani, and Kailin Chen in modifying our HTS Python scripts and HPC execution.

