## Supplemental Figures and Tables for "High throughput screening of mesenchymal stromal cell morphological response to inflammatory signals for bioreactor-based manufacturing of extracellular vesicles that modulate microglia"

### SUPPLEMENTAL FIGURES AND INFORMATION

**Table S1. Cell-line information.**

| Cell-line | Tissue source | Donor age | Donor sex | PDL |
|---|---|---|---|---|
| ADMSC RB62 | Adipose | 31-45 | Female | 15.83 |
| BMMSC RB71 | Bone marrow | 18-30 | Female | 12.72 |
| ADMSC RB98 | Adipose | 18-30 | Female | 11.77 |

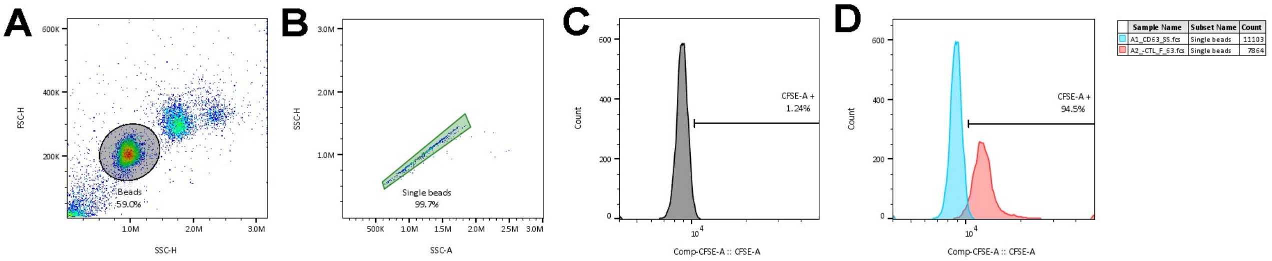


**Figure S1. MSC-EV bead-based flow cytometry gating strategy. A.** Beads were gated based on scatter principles. **B.** Single beads were gated. **C.** CFSE+ was determined by gating CFSE FMO 1% CFSE+. **D.** MSC-EV preparation from -CTL flask batch 1 overlaid on CFSE FMO shows 94.5% CFSE+, indicating intact, CD81+ vesicles.

**Table S2. Metrics to assess MSC-EV preparation quality.**

| Identity | Purity | Potency |
|---|---|---|
| Protein yield (ug/mL) | Protein/particle (ug/particle) | Microglia modulation (composite microglia morphological score) |
| Particle yield (particles/mL) | | |
| Lipid bilayer (CFSE+) | | |
| Canonical surface marker (CD81+) | | |

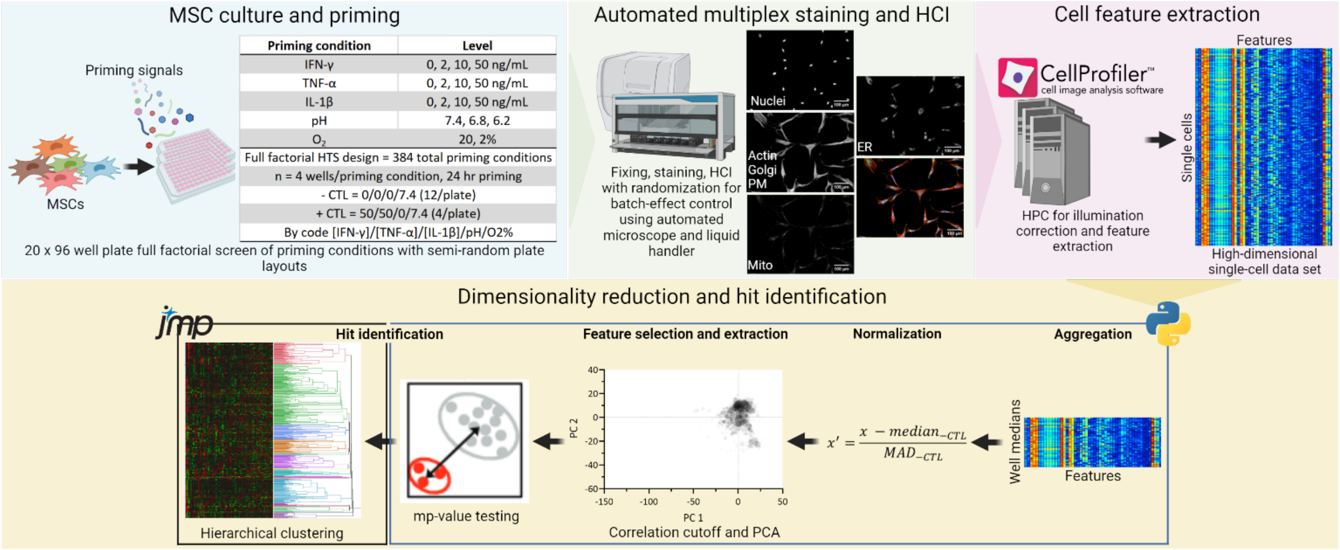


**Figure S2. Exploratory HTS hit priming condition identification workflow.** Created with BioRender.com.

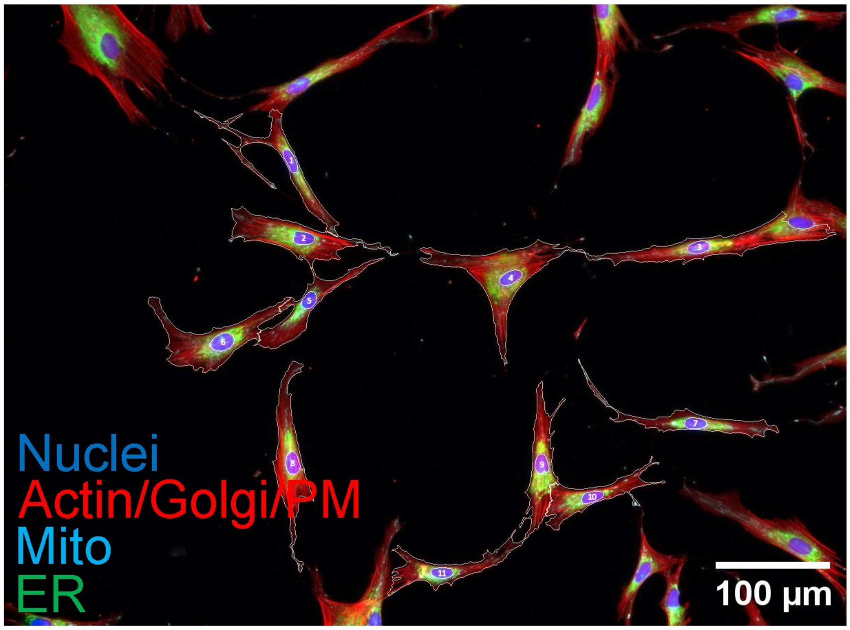


**Figure S3. CellProfiler analysis pipeline segmented nucleus and cell body accurately.** Color composite image with cell body and nucleus segmentation lines (white). Number in nucleus indicates object number in CellProfiler metadata.

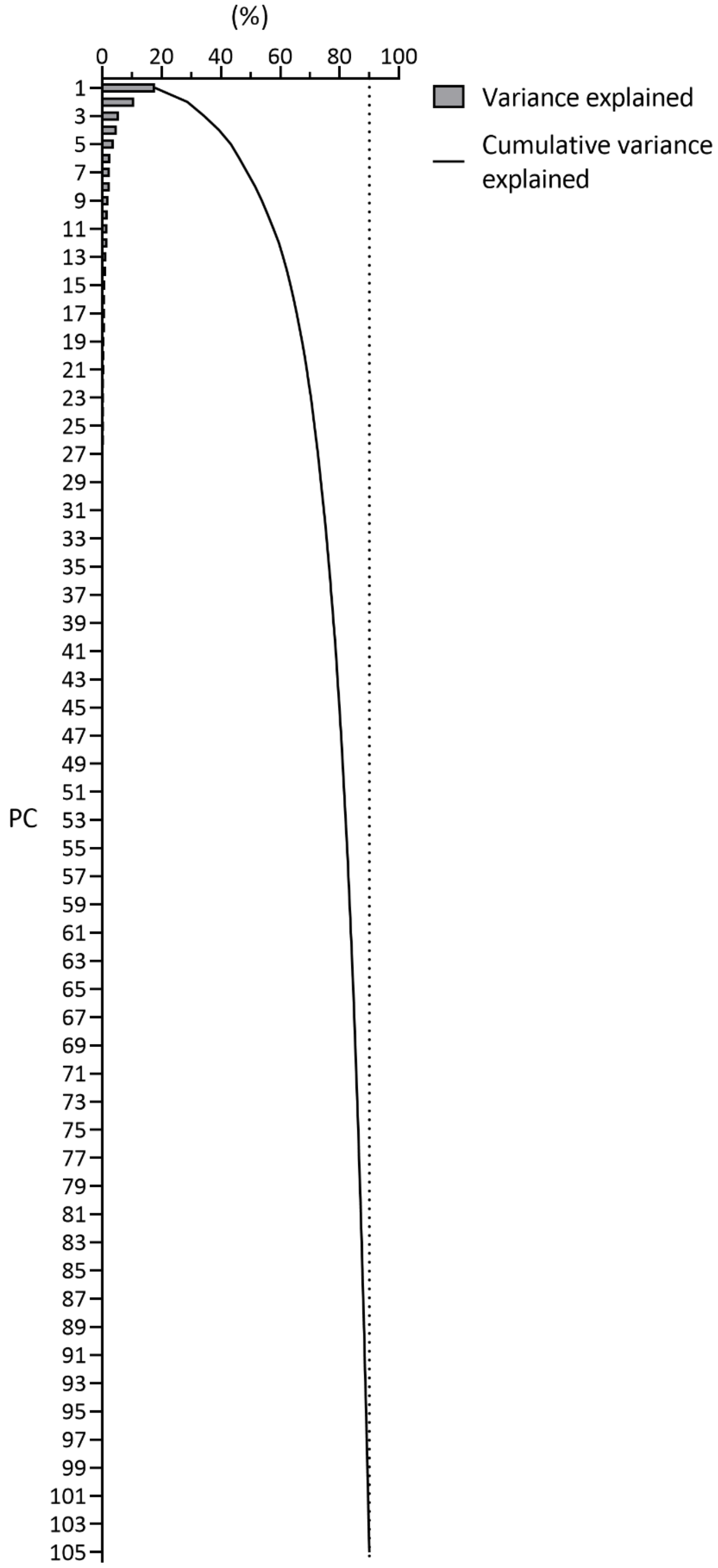


**Figure S4. 105 PCs explained 90% of the variance in the exploratory HTS data.** Scree plot of exploratory HTS PCA.

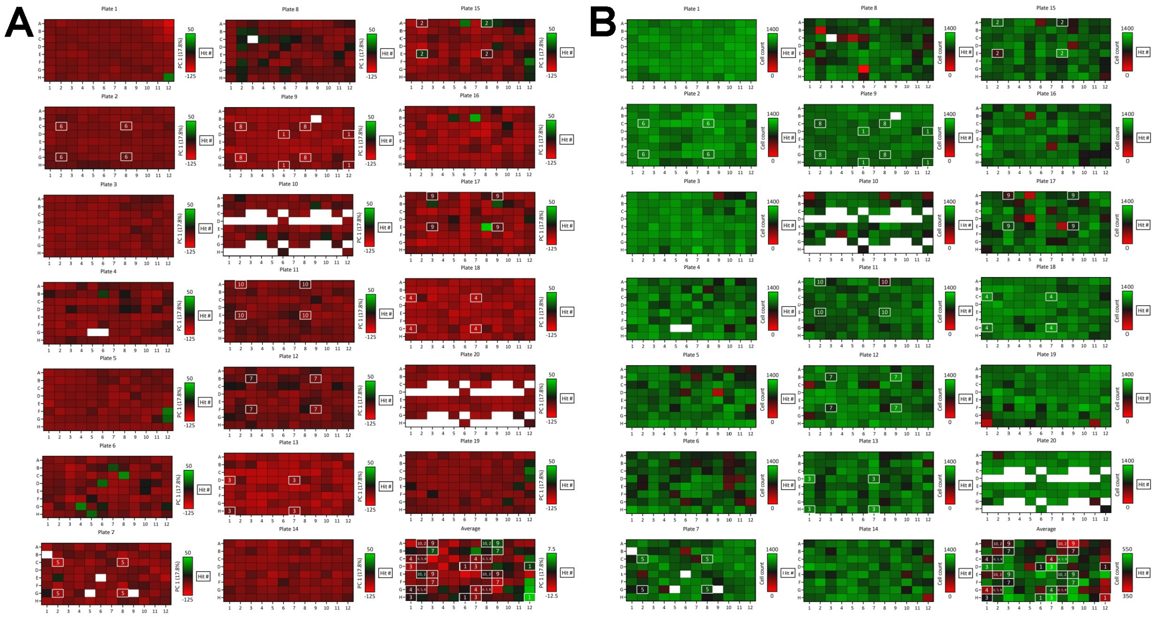

**Figure S5. Visual HTS quality assessment supported that hits were identified based on replicable biological phenomena. A.** Plate heatmap colored by PC1. **B.** Plate heatmap colored by cell count. **A-B.** Plate quadrants are replicate wells. White spaces on plates 10 and 20 are non-experimental wells (filled with RoosterCollect-EV during the experiment); white spaces on plates 4, 7, 8, and 9 are wells intentionally removed from the experiment due to known pipetting error.

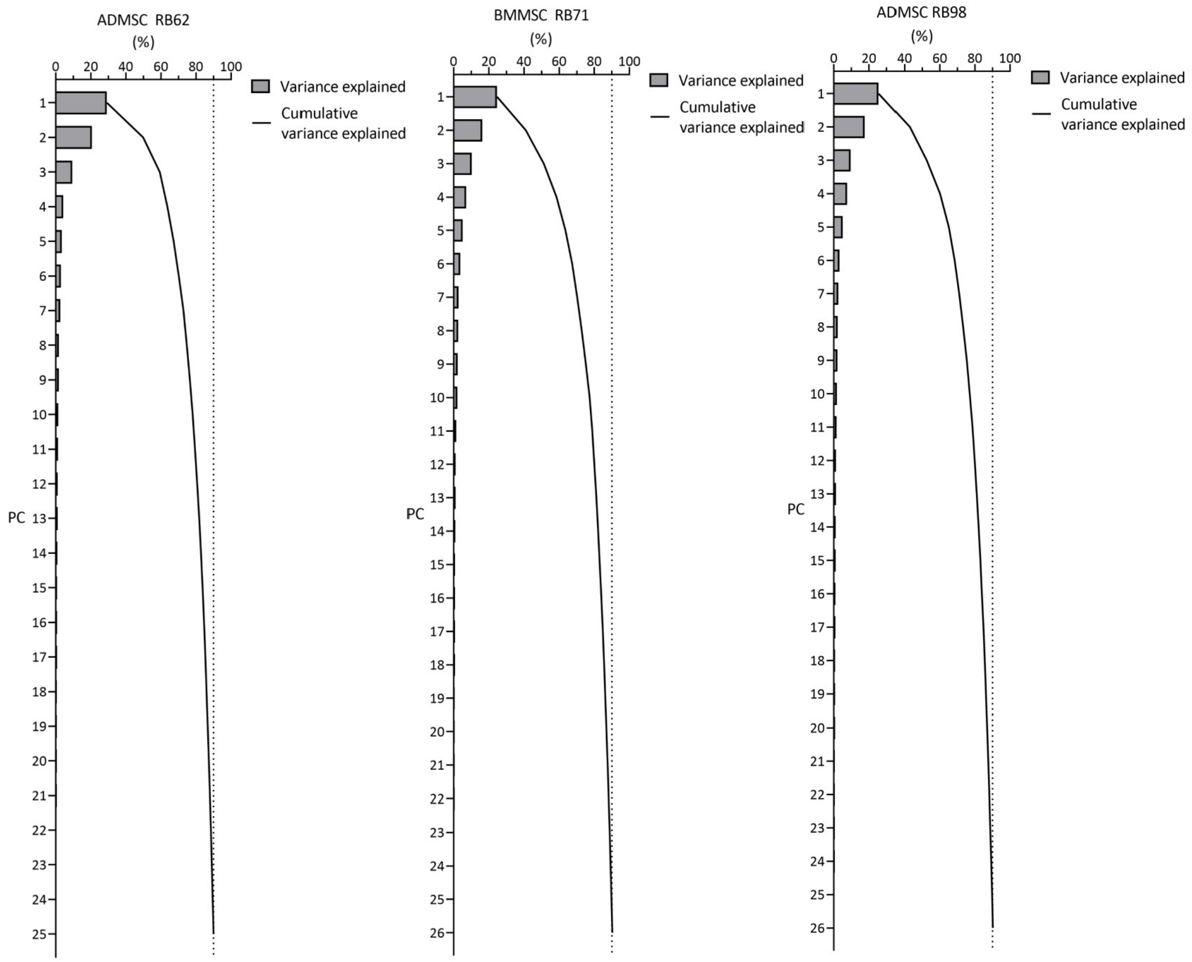


**Figure S6. 26 PCs explained 90% of the variance in the validation HTS data.** Scree plot by cell line of validation HTS PCA.

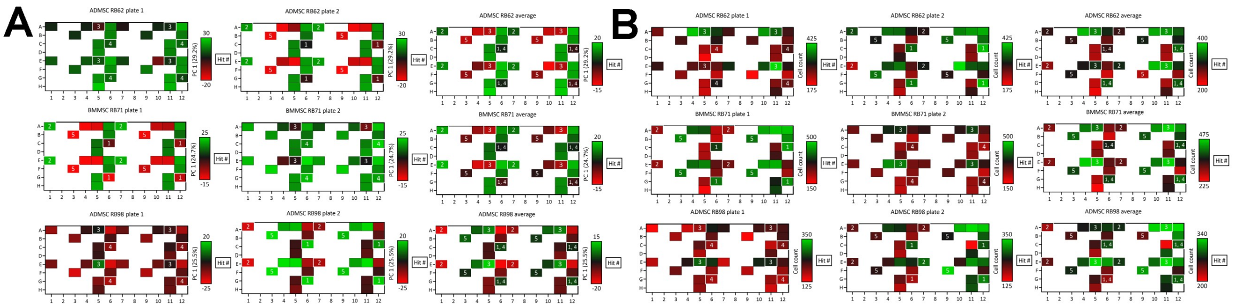

**Figure S7. Visual HTS quality assessment supported that hits were validated based on replicable biological phenomena. A.** Plate heatmap colored by PC1. **B.** Plate heatmap colored by cell count. **A-B.** Plate quadrants are replicate wells. Plate order is randomized but treatment layout is the same for all cell lines. White spaces on all plates are non-experimental wells (filled with RoosterCollect-EV during the experiment).

**Table S3. MSC-EV manufacturing hit priming conditions.**

| Exploratory hit # | Manufacturing hit # |
|---|---|
| 6 | 1 |
| 10 | 2 |
| 4 | 3 |
| 8 | 4 |
| 2 | 5 |

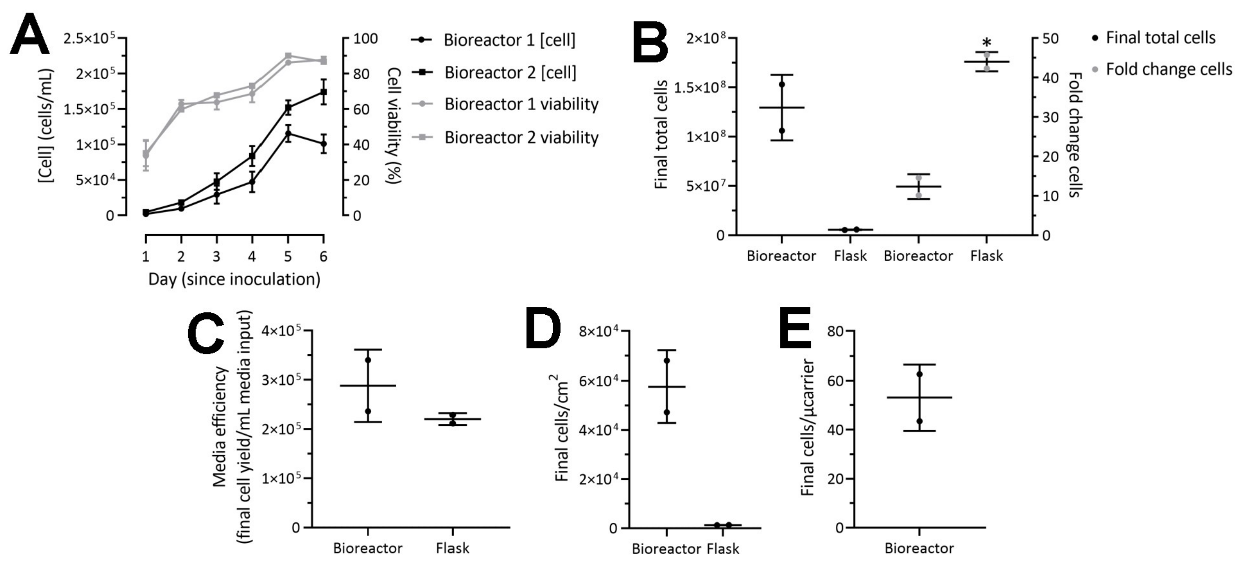


**Figure S8. Bioreactors consistently scaled-up MSC-EV manufacturing. A.** Bioreactor growth curves. Graphed as mean and standard deviation where each point represents 1 sample, n = 3, ns, multiple unpaired t tests with Welch correction. **B-D.** Flask and bioreactor growth characteristics. Graphed as mean and standard deviation where each point represents 1 vessel, n = 2, *p<0.05, unpaired t test with Welch correction. **E.** Bioreactor growth characteristics. Graphed as mean and standard deviation where each point represents 1 vessel, n = 2.

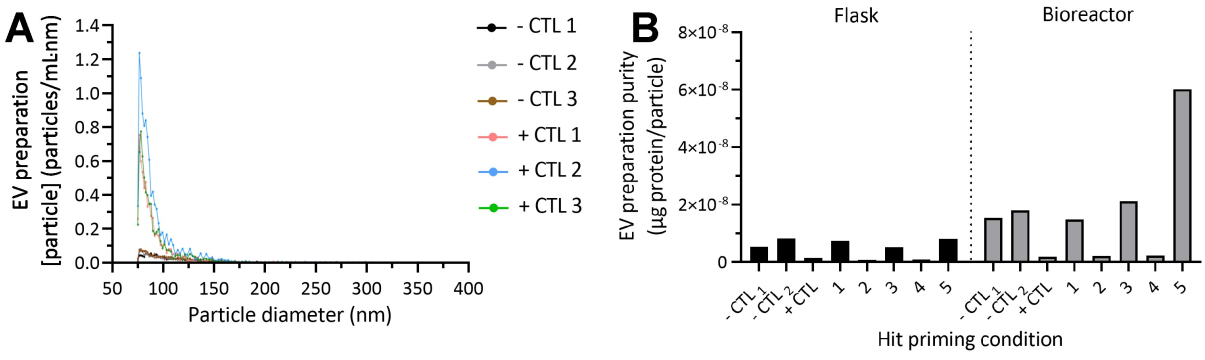


**Figure S9. +CTL, Hit 2, and Hit 4 priming conditions significantly impacted MSC-EV protein and particle yield. A.** Particle count histograms. **D.** Purity from **Fig. 4D** by condition.

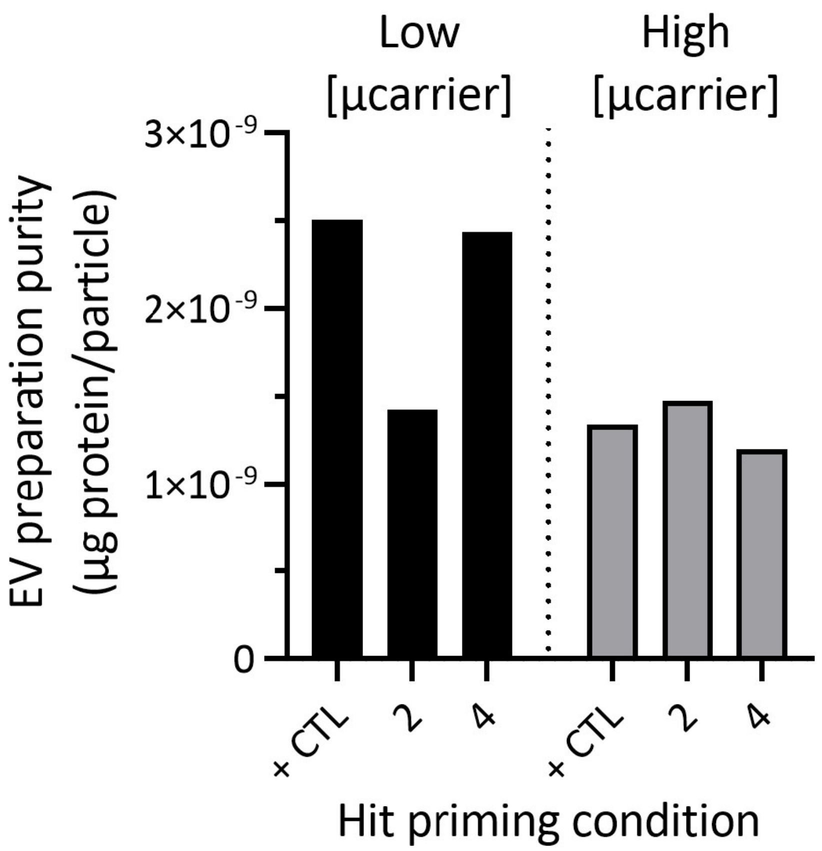


**Figure S10. Validation MSC-EV manufacturing purity by condition.** Same data as plotted in **Fig 6D**.

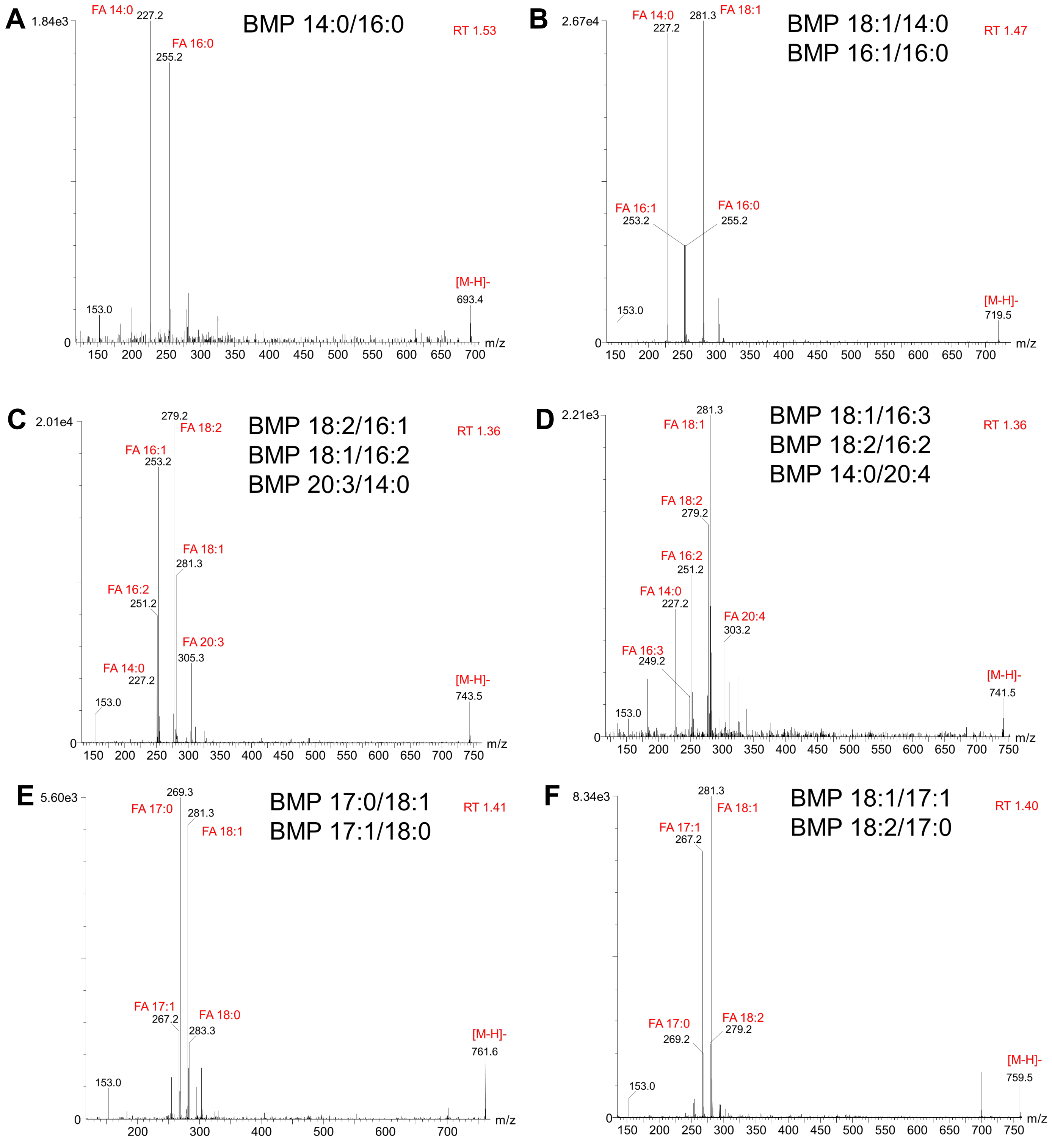

A
1.84e3
BMP 14:0/16:0
RT 1.53
FA 14:0
227.2
FA 16:0
255.2
153.0
[M-H]-
693.4
m/z
B
2.67e4
BMP 18:1/14:0
BMP 16:1/16:0
RT 1.47
FA 14:0
227.2
281.3
FA 18:1
FA 16:1
253.2
FA 16:0
255.2
153.0
[M-H]-
719.5
m/z
C
2.01e4
BMP 18:2/16:1
BMP 18:1/16:2
BMP 20:3/14:0
RT 1.36
279.2
FA 18:2
FA 16:1
253.2
FA 18:1
281.3
FA 16:2
251.2
FA 20:3
305.3
FA 14:0
227.2
153.0
[M-H]-
743.5
m/z
D
2.21e3
BMP 18:1/16:3
BMP 18:2/16:2
BMP 14:0/20:4
RT 1.36
281.3
FA 18:1
FA 18:2
279.2
FA 16:2
251.2
FA 14:0
227.2
FA 20:4
303.2
FA 16:3
249.2
153.0
[M-H]-
741.5
m/z
E
5.60e3
BMP 17:0/18:1
BMP 17:1/18:0
RT 1.41
269.3
FA 17:0
281.3
FA 18:1
FA 17:1
267.2
FA 18:0
283.3
153.0
[M-H]-
761.6
m/z
F
8.34e3
BMP 18:1/17:1
BMP 18:2/17:0
RT 1.40
281.3
FA 18:1
FA 17:1
267.2
FA 17:0
269.2
FA 18:2
279.2
153.0
[M-H]-
759.5
m/z

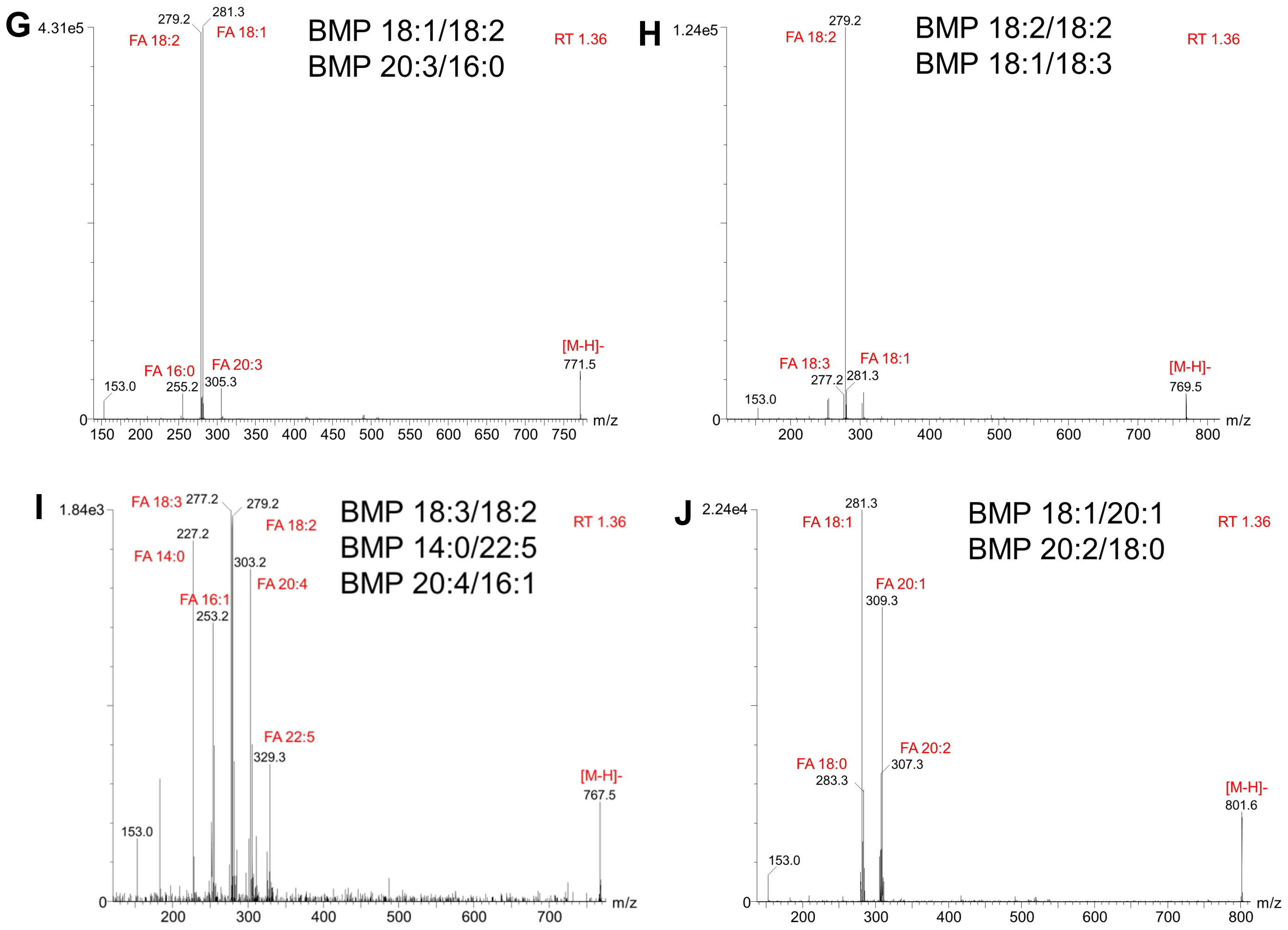


**Figure S11.** MS/MS spectra collected in ESI- with a CE ramp of 45-60 V. The spectra are of the phospholipid class bis(monoacylglycero)phosphate (BMP) with sum compositions of **A)** BMP 30:0 **B)** BMP 32:1 **C)** BMP 34:3 **D)** BMP 34:4 **E)** BMP 35:1 **F)** BMP 35:2 **G)** BMP 36:3 **H)** BMP 36:4 **I)** BMP 36:5 **J)** BMP 38:2. Spectra have been labeled with retention time (RT), precursor [M-H]-, and fatty acid (FA) identities, and isomeric compositions.

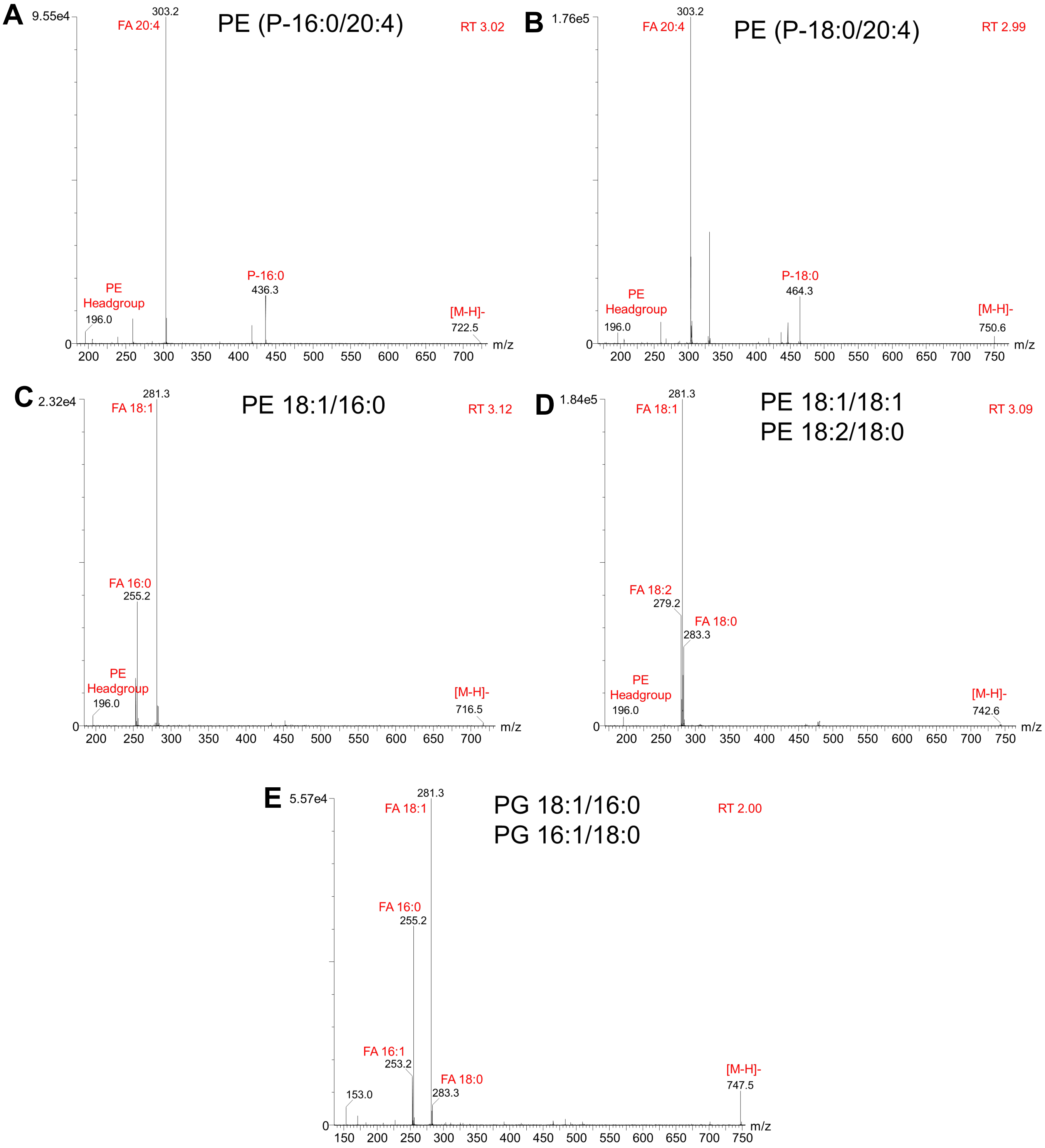


**Figure S12.** MS/MS spectra collected in ESI- with a CE ramp of 45-60 V. The spectra are of the phospholipid classes plasmalogen phosphatidylethanolamine (PE P), phosphatidylethanolamine (PE) and phosphatidylglycerol (PG) with sum compositions of **A)** PE P-36:4 **B)** PE P-38:4 **C)** PE 34:1 **D)** PE 36:2 **E)** PG 34:1**.** Spectra have been labeled with retention time (RT), precursor [M-H]-, and fatty acid (FA) identities, and isomeric compositions.

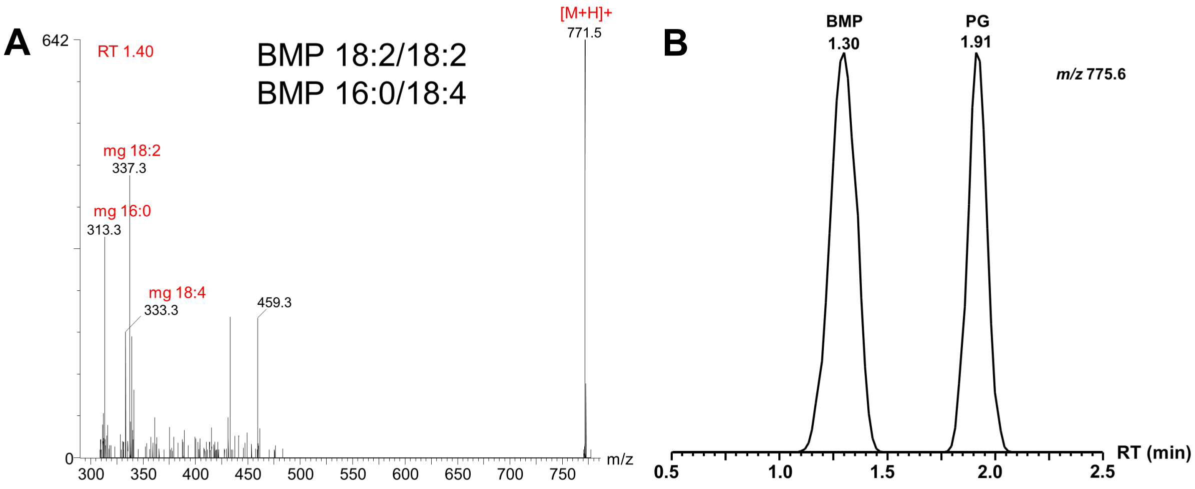


**Figure S13. A)** MS/MS spectrum collected in ESI+ with a CE ramp of 45-60 V. The spectrum is of the phospholipid class bis(monoacylglycero)phosphate (BMP) with sum composition of BMP 36:4. The spectrum has been labeled with retention time (RT), precursor [M+H]+, and monoglyceride [M+H-$H_2O$]+ ion (mg) identities, and isomeric compositions. **B)** Extracted ion chromatograms of BMP 36:1 and PG 36:1 using negative mode electrospray ionization. Despite having identical masses, chromatography completely separates the isomeric classes.

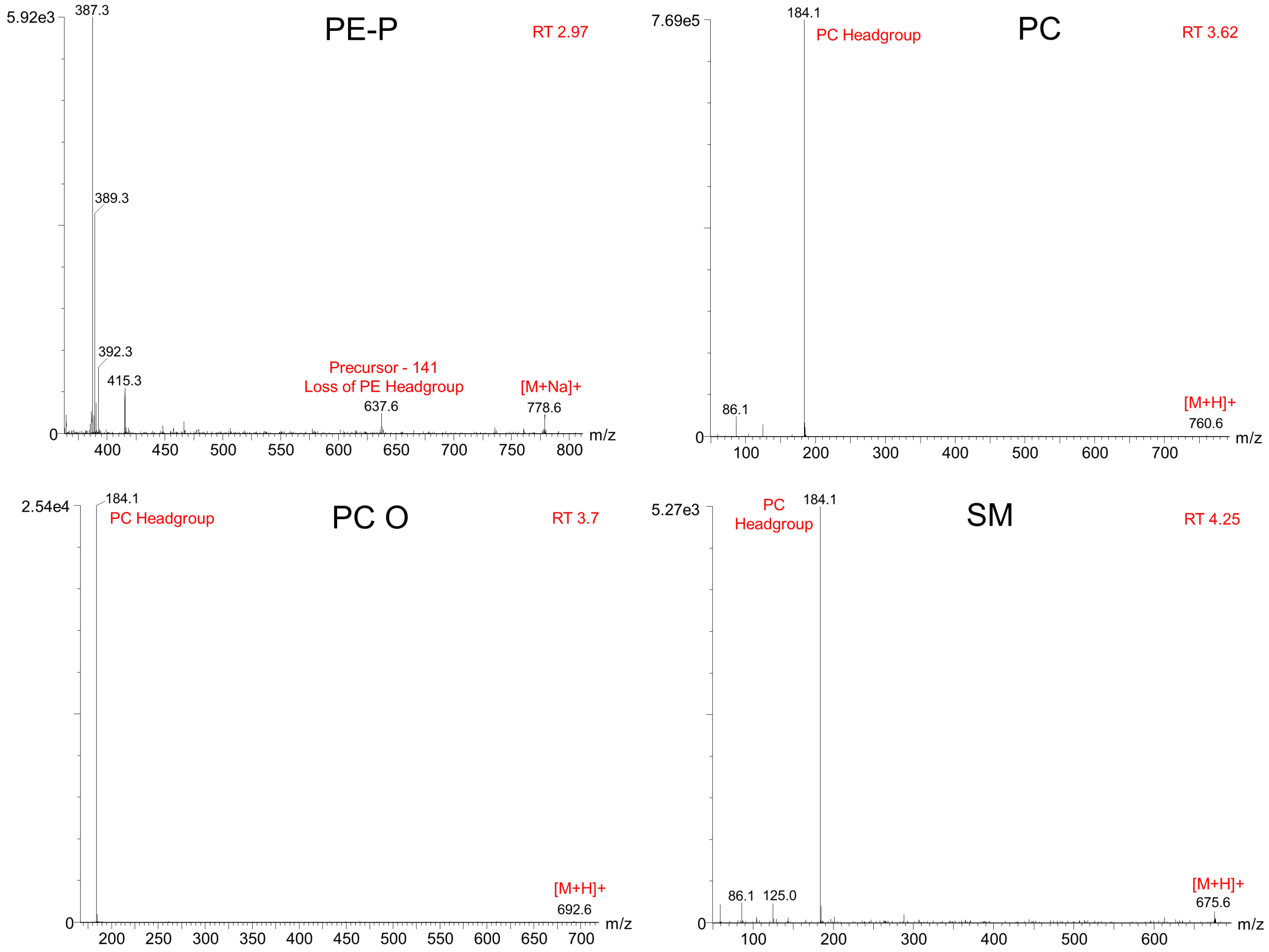


**Figure S14.** MS/MS spectra collected in ESI+ with a CE ramp of 45-60 V. The spectra are representative examples of the phospholipid classes plasmalogen **A)** phosphatidylethanolamine (PE P), **B)** phosphatidylcholine (PC), **C)** phosphatidylcholine with alkyl ether substituent (PC O), and **D)** sphingomyelin (SM). Spectra have been labeled with retention time (RT), precursor [M+H]+ or [M+Na]+, and lipid headgroup information.

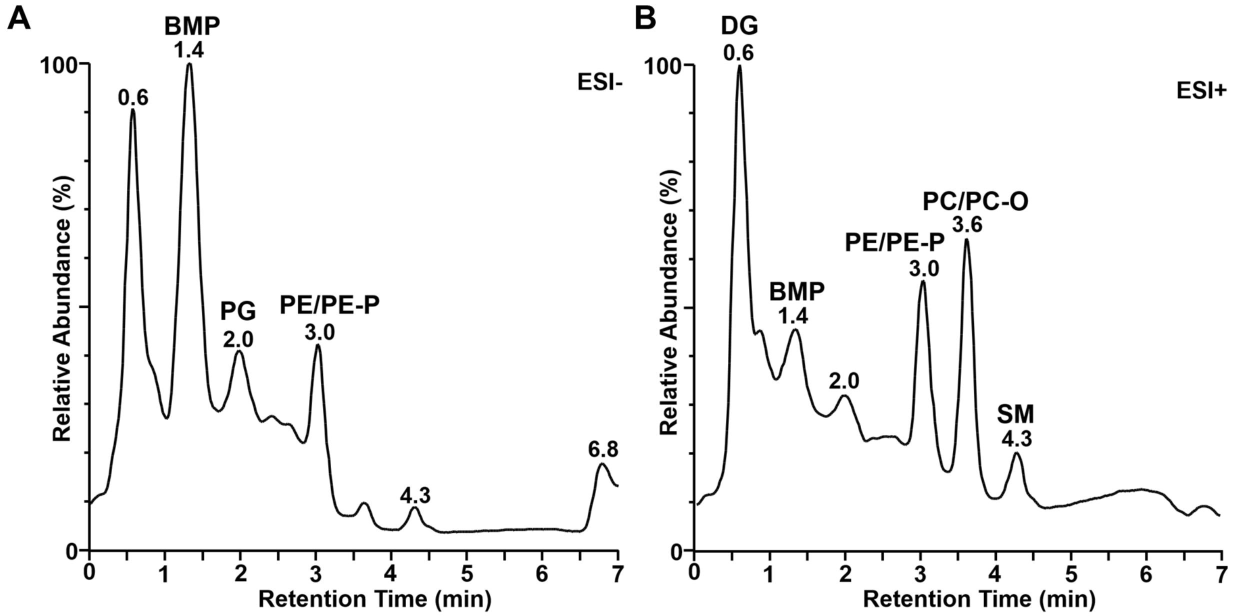


**Figure S15.** Total ion chromatograms of lipids using electrospray ionization in **A)** negative mode and **B)** positive mode. Each lipid class has a characteristic retention time region in which it elutes.
