## Supplementary material for "High throughput screening of mesenchymal stromal cell morphological response to inflammatory signals for bioreactor-based manufacturing of extracellular vesicles that modulate microglia": cytominer-eval_python_script_S6.py

```
#!/usr/bin/env python
# coding: utf-8

# In[1]:


import numpy as np
import pandas as pd
from sklearn import preprocessing

from cytominer_eval import evaluate
from pycytominer.cyto_utils import infer_cp_features


# In[ ]:


#Load Data
df = pd.read_csv("final_data_for_jmp3_no_p21.csv")
print(df.shape)
df = df.dropna(axis=0)
df.head(10)


# In[ ]:


# Make ctrl wells in HS_ng_per_mL column more specific based on bFGF condition
#df.loc[(df["Metadata_bFGF_ng_per_mL"] == 'minus'), "Metadata_HS_ng_per_mL"] = df['Metadata_HS_ng_per_mL'].astype(str) + "_minus"
#df.head(50)


# In[ ]:


#Set Feature Information
features = infer_cp_features(df)
meta_features = infer_cp_features(df,metadata=True)

meta_features


# In[ ]:


control_perts = df.query("Metadata_treatment == '0_0_0_7.4'").Metadata_treatment.unique().tolist()
control_perts


# In[ ]:


mp_values = []
    
    for Treatment in df.Metadata_Treatment.unique():
        Treatment_df = df.query("Metadata_Treatment == @Treatment")
        #print(Treatment_df.head())
    
        for num_perm in [1000]:
    
            mp_results = evaluate(
                profiles=Treatment_df,
                features=features,
                meta_features=meta_features,
                replicate_groups="Metadata_treatment",
                operation="mp_value",
                similarity_metric="pearson",
                mp_value_params={"rescale_pca": True, "nb_permutations": num_perm},
                grit_control_perts=control_perts
            ).assign(Treatment=Treatment, permutations=num_perm)
        
            mp_values.append(mp_results)


# In[ ]:


mp_values = pd.concat(mp_values).reset_index(drop=True)

sig_mp_values = mp_values.loc[mp_values["mp_value"] <= 0.05]

mp_values.to_csv("mp_values_corr0.9_var_3.csv", index=False)
sig_mp_values.to_csv("sig_mp_values_corr0.9_var_iter.csv", index=False)
print(mp_values.shape)
print(sig_mp_values.shape)
sig_mp_values.head()


# In[ ]:


# Group significant mp_value dataframe by similar treatment and HS_ng_per_mL
#sig_mp_values = sig_mp_values.sort_values(by = ['Treatment','Metadata_HS_ng_per_mL'])

# Calculate average mp_value and stdev for each iteration of significant (mp <= 0.05) perturbation
#avg_sig_mp = sig_mp_values.groupby(['Treatment','Metadata_HS_ng_per_mL'],as_index=False)['mp_value'].mean().rename(columns={'mp_value':'avg_mp_value'})
#stdev_sig_mp = sig_mp_values.groupby(['Treatment','Metadata_HS_ng_per_mL'],as_index=False)['mp_value'].std().rename(columns={'mp_value':'stdev_mp_value'})
#sig_mp_values = pd.concat([avg_sig_mp,stdev_sig_mp], axis=1, sort=False)
#sig_mp_values = sig_mp_values.loc[:,~sig_mp_values.columns.duplicated()]

#sig_mp_values.to_csv('avg_sig_mp_value_corr0.9_var1.csv', index=False)
#sig_mp_values.head()
```
