## Supplementary material for "High throughput screening of mesenchymal stromal cell morphological response to inflammatory signals for bioreactor-based manufacturing of extracellular vesicles that modulate microglia": morphology_python_script_S3.py

```
#!/usr/bin/env python
# coding: utf-8

# In[191]:


# Inputs: Cells csv, Filtered Nucleus csv, Experiment Conditions csv
cells_csv_path = '.\Cells_combined.csv'
nucleus_csv_path = r'.\Nucleus_combined.csv'
cytoplasm_csv_path = '.\Cytoplasm_combined.csv'
conditions_csv_path = '.\Conditions.csv'


# In[192]:


import pandas as pd #Python Library for reading and manipulating tabulated data
from scipy.stats import median_abs_deviation


# In[193]:


import numpy as np #Python Library for various operations on the matrices


# In[194]:


# Reading all files  
cells = pd.read_csv(cells_csv_path)
nucleus = pd.read_csv(nucleus_csv_path)
cytoplasm = pd.read_csv(cytoplasm_csv_path)
condition = pd.read_csv(conditions_csv_path)


# In[195]:


#Delete any rows that are copies of the column header
cells = cells.loc[cells["ImageNumber"] != "ImageNumber"]
nucleus = nucleus.loc[nucleus["ImageNumber"] != "ImageNumber"]
cytoplasm = cytoplasm.loc[cytoplasm["ImageNumber"] != "ImageNumber"]
#generate csv when running a test .csv file to see if above code works to delete rows that are just headers
#cells.to_csv('any_objects_left.csv')


# In[196]:


# Processing:cells


# In[197]:


cells.info()
cells.shape


# In[198]:


#Specify datatype for each column -- only Metadata_Well columns should be strings
for column in cells:
    if column == "Metadata_Well" or column == "Metadata_Well.1":
        cells.astype({column: str})
    else:
        #If column contains data, use pd.to_numeric() to decide whether to make datatype equal to "int" or "float"
        cells[column] = pd.to_numeric(cells[column])
cells.info()

for column in nucleus:
    if column == "Metadata_Well" or column == "Metadata_Well.1":
        nucleus.astype({column: str})
    else:
        #If column contains data, use pd.to_numeric() to decide whether to make datatype equal to "int" or "float"
        nucleus[column] = pd.to_numeric(nucleus[column])

for column in cytoplasm:
    if column == "Metadata_Well" or column == "Metadata_Well.1":
        cytoplasm.astype({column: str})
    else:
        #If column contains data, use pd.to_numeric() to decide whether to make datatype equal to "int" or "float"
        cytoplasm[column] = pd.to_numeric(cytoplasm[column], errors='coerce')


# In[199]:


cells.head()


# In[200]:


#### Deletion of unnecessary columns in cells csv


# In[201]:


# columns_to_drop = ['Metadata_ChannelIn',
#         'Metadata_ChannelOut'
#         'Metadata_FileLocation',
#         'Metadata_Frame',
#         'Metadata_Series',
#        'Metadata_Site',
#         'Metadata_Time',
#         'AreaShape_BoundingBoxArea',
#         'AreaShape_BoundingBoxMaximum_X',
#         'AreaShape_BoundingBoxMaximum_Y',
#         'AreaShape_BoundingBoxMinimum_X',
#         'AreaShape_BoundingBoxMinimum_Y',
#         'AreaShape_EulerNumber',
#         'Location_Center_X',
#         'Location_Center_Y',
#         'Location_Center_Z',
#         'Number_Object_Number',
#         'Parent_FilteredNuclei',
#         'Parent_Nuclei',
#         'AreaShape_Orientation']


# In[202]:


# cells.drop(columns_to_drop, axis=1, inplace = True)


# In[203]:


cells.info()
cells.shape


# In[204]:


# Add suffix "cell" to distinguish from nuclei data
cells.columns = [col + '_cell' for col in cells.columns]
nucleus.columns = [col + '_nucleus' for col in nucleus.columns]
cytoplasm.columns = [col + '_cytoplasm' for col in cytoplasm.columns]


# In[205]:


cells.info()


# In[206]:


### Merging Cell and Nucleus


# In[207]:


cellsandnucleus = pd.concat([cells, nucleus, cytoplasm], axis=1, sort=False) 


# In[208]:


cellsandnucleus.info()


# In[209]:


cellsandnucleus.shape


# In[210]:


## Feature Engineering
#### Addition of new data columns based on calculation from existing data


# In[211]:


cellsandnucleus['objectnumbercheck'] = np.where(cellsandnucleus['ObjectNumber_cell'] == cellsandnucleus['ObjectNumber_nucleus'], 0, np.nan)


# In[212]:


cellsandnucleus.info()


# In[213]:


cellsandnucleus.isna().sum() # Ensure no value in the dataframe corresponds to nan = not a number


# In[214]:


cellsandnucleus['NC_Ratio'] = cellsandnucleus['AreaShape_Area_nucleus']/cellsandnucleus['AreaShape_Area_cell']


# In[215]:


cellsandnucleus.info()
cellsandnucleus.shape


# In[216]:


cellsandnucleus['Aspect_Ratio_Cell'] = cellsandnucleus['AreaShape_MajorAxisLength_cell']/cellsandnucleus['AreaShape_MinorAxisLength_cell']


# In[217]:


cellsandnucleus['Aspect_Ratio_Nucleus'] = cellsandnucleus['AreaShape_MajorAxisLength_nucleus']/cellsandnucleus['AreaShape_MinorAxisLength_nucleus']


# In[218]:


cellsandnucleus['PerimeterArea_Ratio_Cell'] = cellsandnucleus['AreaShape_Perimeter_cell']/cellsandnucleus['AreaShape_Area_cell']


# In[219]:


cellsandnucleus['PerimeterArea_Ratio_Nucleus'] = cellsandnucleus['AreaShape_Perimeter_nucleus']/cellsandnucleus['AreaShape_Area_nucleus']


# In[220]:


cellsandnucleus.info()


# In[221]:


cellsandnucleus.isna().sum() 


# In[222]:


cellsandnucleus.head(10)


# In[223]:


cellsandnucleus.shape


# In[224]:


## Data Cleaning based on Engineered Features
### Identifying NC Ratio range and cutoff
#### Weed out poorly segmented cells, the nucleus may be either extremely big or extremely small; NC ratio helps identify the cells with very high or very low nucleus area relative to cell area. NC ratio not = 1 (that would mean the nucleus and cell are the same size). Also this ratio can only be between 0 and 1 
#### Filter NC Ratio area >= 0.75 and delete
#### Plot a distribution of NC Ratio


# In[225]:


import matplotlib.pyplot as plt
def draw_histogram(df, col, x_label, y_label, title):
    ax = df.hist(bins=100, column=col, figsize=(10,5))
    axes = ax.flatten()
    axes[0].set_xlabel(x_label, fontsize=12)
    axes[0].set_ylabel(y_label, fontsize=12)
    axes[0].set_title(title, fontsize =18)


# In[226]:


draw_histogram(df=cellsandnucleus, col='NC_Ratio', x_label='NC_Ratio', y_label='Frequency', title='Distribution of NC Ratio')


# In[227]:


#### Get a more fine-grained count of number of cells for each 0.1 increase in NC ratio


# In[228]:


cellsandnucleus.groupby(pd.cut(cellsandnucleus['NC_Ratio'], np.arange(0, 1.1, 0.1)))['Metadata_Well_cell'].count()


# In[229]:


#### Identify wells where cells with NC Ratio >=0.75 observed, so that they may be visually inspected for poor segmentation


# In[230]:


cellsandnucleus[cellsandnucleus['NC_Ratio'] >= 0.75][['Metadata_Well_cell', 'ObjectNumber_cell', 'NC_Ratio']]


# In[231]:


### Identify outliers based on Nucleus Area


# In[232]:


draw_histogram(df=cellsandnucleus, col='AreaShape_Area_nucleus', x_label='AreaShape_Area_nucleus', y_label='Frequency', title='Distribution of AreaShape_Area_nucleus')


# In[233]:


cellsandnucleus.groupby(pd.cut(cellsandnucleus['AreaShape_Area_nucleus'], np.arange(0, 8500, 500)))['Metadata_Well_cell'].count()


# In[234]:


# cellsandnucleus[(cellsandnucleus['AreaShape_Area_nucleus'] > 2000) & (cellsandnucleus['AreaShape_Area_nucleus'] <= 2200)][['Metadata_Well_cell', 'ObjectNumber_cell', 'AreaShape_Area_nucleus']]


# In[235]:


# cellsandnucleus[(cellsandnucleus['AreaShape_Area_nucleus'] > 2200)]['AreaShape_Area_nucleus'].count()


# In[236]:


cellsandnucleus.drop(cellsandnucleus[cellsandnucleus.NC_Ratio >= 0.75].index, inplace=True)
cellsandnucleus.shape


# In[237]:


# cellsandnucleus.drop(cellsandnucleus[cellsandnucleus.NC_Ratio < 0.05].index, inplace=True)
# cellsandnucleus.shape


# In[238]:


cellsandnucleus.drop(cellsandnucleus[cellsandnucleus.AreaShape_Solidity_nucleus > 1].index, inplace=True)
cellsandnucleus.shape


# In[239]:


cellsandnucleus.drop(cellsandnucleus[cellsandnucleus.AreaShape_Solidity_cell > 1].index, inplace=True)
cellsandnucleus.shape


# In[240]:


# cellsandnucleus.drop(cellsandnucleus[cellsandnucleus.AreaShape_Area_nucleus >= 2200].index, inplace=True)
# cellsandnucleus.shape


# In[241]:


cellsandnucleus.head()


# In[242]:


#cellsandnucleus = cellsandnucleus.loc[:,~cellsandnucleus.columns.duplicated()]


# In[243]:


# calculate well medians

N = cellsandnucleus.groupby(['Metadata_Plate_cell','Metadata_Well_cell'],as_index=False).size()
col_drop = ['Metadata_Plate_cell','Metadata_Well_cell']
N.drop(col_drop,axis=1,inplace=True)
cellsandnucleus = cellsandnucleus.groupby(['Metadata_Plate_cell','Metadata_Well_cell'],as_index=False).median()
cellsandnucleus = pd.concat([N,cellsandnucleus], axis=1, sort=False)


# In[244]:


# (cellsandnucleus['Stimuli'] == 1).sum() # Check for how many cells are stimulated


# In[245]:


cellsandnucleus.head()


# In[246]:


cellsandnucleus.shape


# In[247]:


cellsandnucleus.isna().sum()


# In[248]:


### Add Experimental Condition Data


# In[249]:


cellsandnucleus['Metadata_Plate_cell'] = cellsandnucleus['Metadata_Plate_cell'].apply(str)


# In[250]:


cellsandnucleus['Plate_Well'] = cellsandnucleus['Metadata_Plate_cell'] + cellsandnucleus['Metadata_Well_cell']


# In[251]:


condition.info()


# In[253]:


cellsandnucleus = pd.merge(cellsandnucleus,condition)


# In[ ]:


cellsandnucleus.head()


# In[ ]:


cellsandnucleus.shape


# In[ ]:


cellsandnucleus.isna().sum()


# In[ ]:


cellsandnucleus.to_csv('compiled_data_file_for_jmp.csv', index=False)  


# In[ ]:


## Pixels to Units Conversion (1 pixel = 0.6 um)


# In[ ]:


cellsandnucleus[['AreaShape_Area_cell','AreaShape_Area_nucleus','AreaShape_Area_cytoplasm']] = cellsandnucleus[['AreaShape_Area_cell','AreaShape_Area_nucleus','AreaShape_Area_cytoplasm']].multiply(0.6*0.6, axis='index') 
cellsandnucleus.head(5)


# In[ ]:


cellsandnucleus[['AreaShape_MajorAxisLength_cell',
'AreaShape_MaxFeretDiameter_cell',
'AreaShape_MaximumRadius_cell',
'AreaShape_MeanRadius_cell',
'AreaShape_MedianRadius_cell',
'AreaShape_MinFeretDiameter_cell',
'AreaShape_MinorAxisLength_cell',
'AreaShape_Perimeter_cell',
'AreaShape_MajorAxisLength_nucleus',
'AreaShape_MaxFeretDiameter_nucleus',
'AreaShape_MaximumRadius_nucleus',
'AreaShape_MeanRadius_nucleus',
'AreaShape_MedianRadius_nucleus',
'AreaShape_MinFeretDiameter_nucleus',
'AreaShape_MinorAxisLength_nucleus',
'AreaShape_Perimeter_nucleus',
'AreaShape_MajorAxisLength_cytoplasm',
'AreaShape_MaxFeretDiameter_cytoplasm',
'AreaShape_MaximumRadius_cytoplasm',
'AreaShape_MeanRadius_cytoplasm',
'AreaShape_MedianRadius_cytoplasm',
'AreaShape_MinFeretDiameter_cytoplasm',
'AreaShape_MinorAxisLength_cytoplasm',
'AreaShape_Perimeter_cytoplasm']] = cellsandnucleus[['AreaShape_MajorAxisLength_cell',
'AreaShape_MaxFeretDiameter_cell',
'AreaShape_MaximumRadius_cell',
'AreaShape_MeanRadius_cell',
'AreaShape_MedianRadius_cell',
'AreaShape_MinFeretDiameter_cell',
'AreaShape_MinorAxisLength_cell',
'AreaShape_Perimeter_cell',
'AreaShape_MajorAxisLength_nucleus',
'AreaShape_MaxFeretDiameter_nucleus',
'AreaShape_MaximumRadius_nucleus',
'AreaShape_MeanRadius_nucleus',
'AreaShape_MedianRadius_nucleus',
'AreaShape_MinFeretDiameter_nucleus',
'AreaShape_MinorAxisLength_nucleus',
'AreaShape_Perimeter_nucleus',
'AreaShape_MajorAxisLength_cytoplasm',
'AreaShape_MaxFeretDiameter_cytoplasm',
'AreaShape_MaximumRadius_cytoplasm',
'AreaShape_MeanRadius_cytoplasm',
'AreaShape_MedianRadius_cytoplasm',
'AreaShape_MinFeretDiameter_cytoplasm',
'AreaShape_MinorAxisLength_cytoplasm',
'AreaShape_Perimeter_cytoplasm']].multiply(0.6, axis = 'index')    


# In[ ]:


cellsandnucleus[['PerimeterArea_Ratio_Cell']] = cellsandnucleus[['PerimeterArea_Ratio_Cell']].multiply(1/0.6, axis='index') 
cellsandnucleus.head(5)


# In[ ]:


cellsandnucleus[['PerimeterArea_Ratio_Nucleus']] = cellsandnucleus[['PerimeterArea_Ratio_Nucleus']].multiply(1/0.6, axis='index') 
cellsandnucleus.head(5)


# In[ ]:


cellsandnucleus.info()


# In[ ]:


cellsandnucleus.head()


# In[ ]:


# cellsandnucleus[cellsandnucleus['Metadata_Well_cell'] == 'F2']


# In[ ]:


## Data Visualization
#### Find outliers based on area, perimeter of cell


# In[ ]:


# draw_histogram(df=cellsandnucleus, col='AreaShape_Area_cell', x_label='Cell Area in um^2',
             # y_label='Frequency', title='Distribution of Cell Area')


# In[ ]:


# max_area = cellsandnucleus['AreaShape_Area_cell'].max()
# min_area = cellsandnucleus['AreaShape_Area_cell'].min()
# print(f'Max area = {max_area} and min area = {min_area}')


# In[ ]:


#### Assess outliers


# In[ ]:


# cellsandnucleus.groupby(pd.cut(cellsandnucleus['AreaShape_Area_cell'], np.arange(0, 15000, 500)))['AreaShape_Area_cell'].count()


# In[ ]:


# cellsandnucleus[((cellsandnucleus['AreaShape_Area_cell'] >= 4000) & (cellsandnucleus['AreaShape_Area_cell'] <= 4500))]
# cellsandnucleus[(cellsandnucleus['AreaShape_Area_cell'] >= 4000)]


# In[ ]:


# cellsandnucleus[(cellsandnucleus['ObjectNumber_cell'] == 1937) & (cellsandnucleus['Metadata_Well_cell'] == 'F2')]['NC_Ratio'] 


# In[ ]:


#### Assess filtering


# In[ ]:


# cellsandnucleus.drop(cellsandnucleus[cellsandnucleus.AreaShape_Area_cell >= 6300].index, inplace=True)
# cellsandnucleus.shape


# In[ ]:


cellsandnucleus.to_csv('compiled_data_file_units.csv', index=False) 
cellsandnucleus.shape


# In[ ]:


#cellsandnucleus.drop(['Metadata_Well_cell',
#                      'Metadata_ChannelIn_cell',
#                      'Metadata_ChannelOut_cell',
#                      'Metadata_FileLocation_cell',
#                      'Metadata_Frame_cell',
#                      'Metadata_Plate_cell',
 #                     'Metadata_Series_cell',
  #                    'Metadata_Site_cell',
   #                   'Metadata_Time_cell',
    #                  'Number_Object_Number_cell',
     #                 'Parent_FilteredNuclei_cell',
      #                'Parent_Nuclei_cell',
       #               'ImageNumber_nucleus',
        #              'ObjectNumber_nucleus',
         #             'Metadata_ChannelIn_nucleus',
          #            'Metadata_ChannelOut_nucleus',
           #           'Metadata_FileLocation_nucleus',
            #          'Metadata_Frame_nucleus',
             #         'Metadata_Plate_nucleus',
              #        'Metadata_Series_nucleus',
#                      'Metadata_Site_nucleus',
 #                     'Metadata_Time_nucleus',
  #                    'Number_Object_Number_nucleus',
   #                   'Parent_Nuclei_nucleus',
    #                  'objectnumbercheck',
     #                 'Plate_Well',
      #                'Metadata_Well',
       #               'Metadata_Plate',
        #              'Metadata_Plate_Well',
         #             'Treatment',
          #            'Metadata_Well_nucleus'
# ], axis=1, inplace = True)


# In[ ]:


## Normalization


# In[ ]:


#from sklearn import preprocessing 
#def get_min_max_scaler(col_vals):
    #min_max_scaler = preprocessing.MinMaxScaler()
    #min_max_scaler.fit(col_vals)
    #return min_max_scaler


# In[ ]:


#dtypes = set()
#scaler_dict = {}
#for col in cellsandnucleus.columns:
    #col_dtype = cellsandnucleus[col].dtype
    #if col_dtype == 'float64' :
        #print(f'Normalizing column {col}')
        #scaler_dict[col] = get_min_max_scaler(cellsandnucleus[col].values.reshape(-1,1))


# In[ ]:


#for col in cellsandnucleus.columns:
    #col_dtype = cellsandnucleus[col].dtype
    #if col_dtype == 'object':
        #print(f'{col}')


# In[ ]:


#for k, scaler in scaler_dict.items():
    #print(f'COL {k} min {scaler.data_min_} max {scaler.data_max_} diff {scaler.data_range_} num_samples {scaler.n_samples_seen_}')


# In[ ]:


#for k, scaler in scaler_dict.items():
    #cellsandnucleus[k] = scaler.transform(cellsandnucleus[k].values.reshape(-1,1))


# In[ ]:


#cellsandnucleus.head()


# In[ ]:


#cellsandnucleus.shape


# In[ ]:


#cellsandnucleus.to_csv('compiled_data_file_normalized.csv')


# In[ ]:


#data = []
#columns = ['Feature', 'Min', 'Max', 'Range', 'Sample_Count']
#for k, scaler in scaler_dict.items():
    #data.append([k, scaler.data_min_[0], scaler.data_max_[0], scaler.data_range_[0], scaler.n_samples_seen_])
#pd.DataFrame(data=data, columns=columns).to_csv('compiled_data_file_normalization_stats.csv', index=False)


# In[ ]:


for plate_num in cellsandnucleus['Metadata_Plate'].unique():
    print(cellsandnucleus[cellsandnucleus['Metadata_Plate'] == plate_num].shape)


# In[ ]:


cellsandnucleus['Treatment'].unique()


# In[ ]:


cellsandnucleus['[bFGF] (ng/mL)'].unique()


# In[ ]:


# Just investigating the number of positive control rows in all plates
cellsandnucleus.loc[(cellsandnucleus['Treatment'] == 'rHS 02 + 100 ng/mL bFGF') & (cellsandnucleus['Concentration rHS (ng/mL)'] == '50000')].shape


# In[ ]:


# Just investigating the number of negative control rows in all plates
cellsandnucleus.loc[(cellsandnucleus['[rHS] (ng/mL)'] == 'ctrl') & (cellsandnucleus['[bFGF] (ng/mL)'] == '0')].shape


# In[ ]:


##Control Normalization


# In[ ]:


# Create a dictionary (plates) which looks like this : {'1': plate1, '2': plate2}. plate1 and plate2 contain rows corresponding to plates 1 and 2 only respectively
plates = {}
# Iterate over each unique plate in the csv
for plate_num in cellsandnucleus['Metadata_Plate'].unique():
    plates[plate_num] = cellsandnucleus[cellsandnucleus['Metadata_Plate'] == plate_num]
    print(f'Identified plate {plate_num} with shape {plates[plate_num].shape}')


# In[ ]:


# Print all non-numeric columns
print('NON-NUMERIC COLUMNS:')
print('--------------------')
for i in cellsandnucleus.columns:
    if cellsandnucleus[i].dtype not in ('float64', 'int64') :
        print(i, cellsandnucleus[i].dtype)


# In[ ]:


normalized_plates = {}
positive_control_means = {} # This would look like { 1: {col1: mean, col2: mean}, 2: {col1:mean, col2: mean}}
negative_control_medians = {}  # This would look like {1: {col1: mean, col2: mean}, 2: {col1:mean, col2: mean}}
negative_control_mads = {}


# In[ ]:


for plate_num, plate in plates.items():
    print(f'PLATE {plate_num}: Processing plate.')
    # Initializing various data structures
    normalized_plate_columns = {}
    #positive_control_means[plate_num] = {}
    negative_control_medians[plate_num] = {}
    negative_control_mads[plate_num] = {}
    
    # Get all rows in this plate where Treatment is 'rHS 02 + 100 ng/mL bFGF' AND 'Concentration rHS (ng/mL)' is 50000
    positive_control_rows = plate.loc[(plate['Treatment'] == 'rHS 02 + 100 ng/mL bFGF') 
                                              & (plate['Concentration rHS (ng/mL)'] == '50000')]
    
    negative_control_rows = plate.loc[(plate['[rHS] (ng/mL)'] == 'ctrl') 
                                              & (plate['[bFGF] (ng/mL)'] == '0')]
    print(f'PLATE {plate_num}: Shape of positive control rows: {positive_control_rows.shape}')
    print(f'PLATE {plate_num}: Shape of negative control rows: {negative_control_rows.shape}')
    
    for col in plate.columns:
        col_dtype = plate[col].dtype
        if col_dtype in ('float64', 'int64'): 
            print(f'PLATE {plate_num}: Normalizing column {col}')
            # Calculate mean of positive control for this col in this plate
            #positive_control_mean = positive_control_rows[col].mean()
            #positive_control_means[plate_num][col] = positive_control_mean
            
            # Calculate median of negative control for this col in this plate
            
            negative_control_median = negative_control_rows[col].median()
            negative_control_medians[plate_num][col] = negative_control_median
            
            negative_control_mad = median_abs_deviation(negative_control_rows[col])
            negative_control_mads[plate_num][col] = negative_control_mad
            
            
            #Subtract the median from each datapoint in the column 
            
            
            # Normalize
            normalized_plate_columns[col] = plate[col].apply(lambda x: (x - negative_control_median)/(negative_control_mad))
        else:
            print (f'PLATE {plate_num}: Skipping normalization of {col} {col_dtype}')
            normalized_plate_columns[col] = plate[col]
        
    # Save the normalized plate in a dictionary
    normalized_plates[plate_num] = pd.concat(normalized_plate_columns, axis=1)
    print('------------------------------------------')


# In[ ]:


## Post-Normalization Validation


# In[ ]:


#col_to_inspect = 'Texture_Variance_OrigDAPI_10_03_256_cytoplasm'
#plate_to_inspect = 1
#pos_mean = positive_control_means[plate_to_inspect][col_to_inspect]
#neg_mean = negative_control_means[plate_to_inspect][col_to_inspect]


# In[ ]:


#normalized_plates[1]['Texture_Variance_OrigDAPI_10_03_256_cytoplasm']


# In[ ]:


#plates[1]['Texture_Variance_OrigDAPI_10_03_256_cytoplasm']


# In[ ]:


#(360.416667 - neg_mean) / (pos_mean - neg_mean)


# In[ ]:


#Save normalization to disk
final_normalized_plate = pd.concat([df for df in normalized_plates.values()])


# In[ ]:


final_normalized_plate.shape


# In[ ]:


final_normalized_plate.to_csv('normalized_plate_combined.csv', index=False)
```
